# Regioisomer-controlled red-shifted DNA probes for imaging of living tissues

**DOI:** 10.64898/2025.12.11.693647

**Authors:** Kamila A. Kiszka, Shalini Pradhan, Jonas Bucevičius, Tanja Koenen, Gražvydas Lukinavičius

**Author notes:** Corresponding author. Email address (G. Lukinavičius). These authors contributed equally to this work (Kamila A. Kiszka, Shalini Pradhan).

## Abstract

Fluorescence imaging of chromatin DNA with high resolution and specificity is key to understanding cellular processes and enabling molecular diagnostics. However, choosing the best DNA probe for *in vivo* imaging is often a challenging task, as systematic studies investigating the biocompatibility of these molecules are lacking. Red-shifted fluorescent probes are particularly advantageous for imaging thick tissues, as light scattering decreases at red and far-red wavelengths. We synthesized and characterized the 4′, 5′, and 6′ carboxyrhodamine regioisomers of SiR-Hoechst and 610CP-Hoechst DNA probes and assessed their performance in mice following intravenous injection. All variants entered tissues and produced stable staining that persisted for several hours *ex vivo*. In particular, 5′ regioisomers show the highest performance, yielding bright and specific nuclear labeling. Renal clearance of the probes was nearly complete within 24 h, as indicated by fluorescence analysis of urine samples. Owing to their low cytotoxicity, high specificity, and favourable photophysical properties, these probes enabled high-quality confocal, two-photon, and STED microscopy in liver, kidney, lung, and heart tissue. Our findings extend the application of rhodamine-based DNA probes to *in vivo* imaging highlight their potential for deep-tissue imaging in live animals.

## 1. Introduction

Organization of chromatin plays a crucial role in gene expression regulation, propagation and preservation of genetic information. Pathological processes can change the arrangement of genes and global chromatin condensation. Nuclear shape abnormalities have been linked to cancer and used in its diagnosis since the 19th century. Modern pathology evaluates metrics like nuclear size, contour irregularities, hyperchromasia, and chromatin distribution to grade cancer ^[1]^. Furthermore, DNA stains are used for infectious diseases diagnostics, such as malaria or trypanosomiasis ^[2]^. Thus, DNA staining dyes are fundamental in various diagnostic applications, from cancer detection to infectious disease identification and genetic disorder screening.

Despite significant advancements in DNA staining for chromatin research and disease diagnostics, several challenges persist. Issues, such as dye toxicity and photobleaching, along with the need for highly specific and sensitive staining protocols, necessitate continuous improvement and innovation in DNA staining techniques. The development of novel dyes and staining methods that offer enhanced specificity, stability, and safety is crucial for advancing diagnostic capabilities ^[3]^. We focused our attention on Hoechst derivatives which emit in red or far-red spectral region ^[4]^. Previously, we analyzed the performance of these DNA stains in cultured living cells and extracted living tissues ^[4a, 5]^. These fluorescent dyes demonstrate low toxicity, high resistance to photobleaching, high specificity and excellent cell-membrane permeability. However, the performance of these stains in living animals remained to be tested.

To this end, we selected a mouse model to perform such experiments and investigate the behavior of the Hoechst-dye conjugates within complex environment of a living animal. In particular, 610CP- and SiR-Hoechst dyes were injected into bloodstream via the tail vein of the mouse at a dosage of 10 mg/kg of body weight. Three positional isomers of the linker attachment point to lower benzene ring (at the 4-,5- and 6-position) of each probe were analyzed. We examined staining of lung, liver, heart, kidney and brain tissues after *post mortem* extraction of the organs. Across all examined tissues, except the brain, staining with 5’ regioisomers produced the highest-quality single- and two-photon fluorescence microscopy images. Furthermore, we demonstrate that upon injection, the 610CP-Hoechst and SiR-Hoechst probes interact with serum albumin, which likely contributes to the retention of Hoechst probes in the body. Analysis of urine samples indicated that the probes were mostly excreted after 24 h post-injection. This reflects the efficient elimination of unbound probe, which further enhances the contrast in the imaging experiment.

In summary, we identified 5-610CP-Hoechst and 5-SiR-Hoechst as fluorescent DNA stains suitable for DNA staining in living animals. These probes show strong potential to serve as diagnostic tools for chromosomal abnormalities, cancer, and infectious diseases.

## 2. Results

### 2.1. Chemical synthesis and photophysical properties of fluorescent DNA probes

We set out to evaluate the *in vivo* performance of positional isomers of red-shifted DNA probes in mice. In our previous work, we reported the synthesis and application of the 5’ and 6’ regioisomers of 610CP and SiR DNA probes (Figure 1a)^[4a]^. More recently, we introduced the 4’ regioisomers of rhodamines, prompting us to investigate their performance in living mice along with previously described 5’ and 6’ regioisomers (Figure 1a). To this end, we prepared fluorescent DNA probes incorporating the 4’ regioisomers of 610CP or SiR dyes by coupling the previously described Hoechst-C4-NH derivative with the corresponding dye carboxylates (4-610CP-COOH or 4-SiR-COOH) using hexafluorophosphate azabenzotriazole tetramethyl uronium (HATU) ^[4a]^. This straightforward synthetic route afforded high yields (60–70%), providing sufficient quantities of fluorescent DNA probes for animal studies **(Figure S1**).

**Figure 1.**
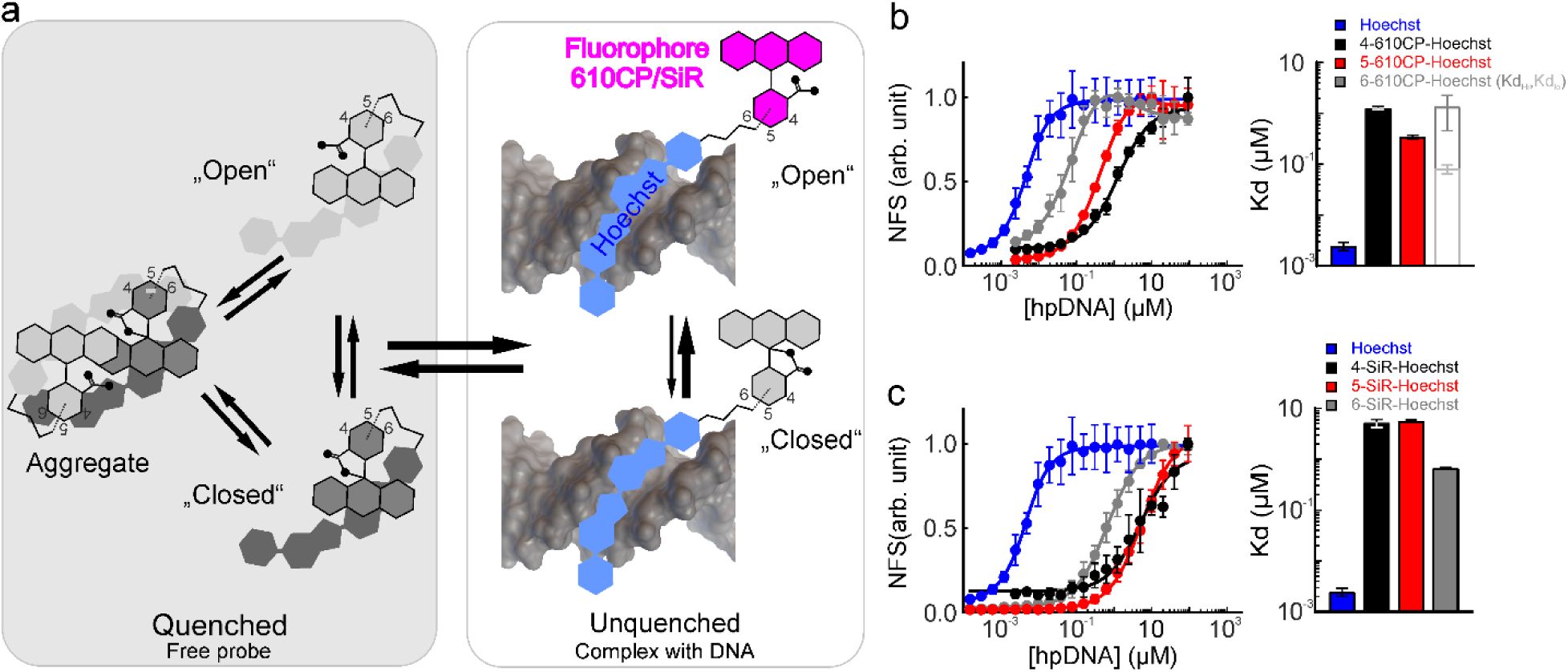
Interaction between Hoechst rhodamine derivatives and hairpin DNA. **a** Proposed model of Hoechst rhodamine conjugates interaction with the target DNA. Free probe is kept in the dark state via three mechanisms – spirolactone formation, intramolecular quenching and aggregation. Minor groove binding results in a fluorescent complex due to equilibrium shifted towards “open” (zwitterion) form, unquenching and disaggregation. **b** Titration of 4nM Hoechst 33342, 3nM 4-610CP-Hoechst, 10nM 5-, 6-610CP-Hoechst with hpDNA. **c** Titration of 4nM Hoechst 33342, 3nM 4-SiR-Hoechst, 10nM 5-, 6-SiR-Hoechst with hpDNA. Two site-binding equation is fitted to the data points for 6-610CP-Hoechst. A single site binding equation was fitted to the data for other regioisomers. All data points are presented as mean ± SD, *n*≥ 3 independent measurements. Apparent Kd values are presented as mean ± SEM. NFS, normalized fluorescence intensity.

Previously, we have observed dramatic quantum yield (QY) decrease of 610CP/SiR 5’/6’ regioisomers after conjugation to Hoechst. The newly synthesized 4’ regioisomer’s probes behaved in the same way and displayed similar low quantum yields in PBS (**Table S1**). As previously for 5’ and 6’ regioisomers, we hypothesize that it is a consequence of self-aggregation and intramolecular quenching. Indeed, addition of 0.1% SDS recovers both, the absorbance and the fluorescence, which is consistent with dissolving aggregates (**Table S1** and **Figure S2**). Thus, the main mechanism of fluorescence quenching is by intermolecular aggregation and π–π interaction (**Figure 1a**).

### 2.2. Interaction of fluorescent DNA probes with the target hpDNA

We have already reported the measurements of the interaction between the target hpDNA and 5/6-610CP-Hoechst or 5/6-SiR-Hoechst probes ^[4a]^. Thus, in this study, we evaluated the interaction of 610CP/SiR 4’ regioisomer probes with the target hpDNA, which resulted in a significant increase in fluorescence and allowed measurement of the K_d_ (**Figure 1b, c** and **Figure S2**). The obtained data points fitted to a single site binding equation, and the dissociation constants (Kd = 1.3 ± 0.1 µM for 4-610CP-Hoechst and Kd = 5.2 ± 0.9 µM for 4-SiR-Hoechst) were similar to previous obtained for 5’/6’ regioisomer probes (**Figure 1 b,c** and **Table S2**). However, the observed fluorescence of the complex with hpDNA for 4’ regioisomer probes was darker compared to the corresponding probes of 5’/6’ regioisomer (**Figure S2** and **Table S2**). This might be due to stronger quenching by DNA heterocyclic bases, resulting from the significantly different conformation of rhodamine 4’ regioisomers compared to the other two regioisomers. The measured quantum yield (QY) of the probe-hpDNA complex for 4’ regioisomers was intermediate between the high value obtained for 5’ regioisomer and the low value for 6’ regioisomer. Interestingly, fluorescence lifetime measurements revealed a single exponential decay for both 4-610CP-Hoechst and 4-SiR-Hoechst probes bound to the target hpDNA (**Table S2**).

### 2.3. Interaction of Hoechst probes with serum albumin

Injection of the fluorescent DNA probes into the bloodstream is likely to result in complex formation between the relatively hydrophobic dye and serum albumin, the most abundant blood protein^[6]^. Indeed, significant increases in absorbance and fluorescence were observed after the addition of high concentrations of mouse, human or bovine serum albumin to the solutions of Hoechst-based DNA probes (**Figure 2a,c** and **Figure S3**). This suggests that the complex formation with serum albumin results in spirolactone opening, and diminished quenching induced by intramolecular *π*-*π* interaction and aggregation. We used this property to determine apparent dissociation constants of the DNA probe-serum albumin complexes (**Figure 2a-d** and **Tables S4-S6**). The binding data for 4/6-610CP and 4/6-SiR probes could be fitted to a single site binding model with apparent K_d_s for mouse serum albumin (MSA) in the range of 10–100 µM (**Figure S4** and **Table S3**). In contrast, the data points of 5-SiR-Hoechst probe fit to two-site binding equation with the first apparent K_d_ located in 10-100 µM range and the second apparent K_d_ exceeding 1000 µM (outside physiological range). Unlike the primary binding event, the secondary binding event resulted in a fluorescence decrease. We hypothesized that similar interaction could take place with human and bovine serum albumins. Titration of human serum albumin (HSA) yielded similar behaviour for both, 5’ and 6’ regioisomer DNA probes (**Figure S4** and **Table S4**). In contrast, bovine serum albumin (BSA) demonstrated single site binding for all positional isomers of both 610CP and SiR fluorophores (**Figure S4** and **Table S5**).

**Figure 2.**
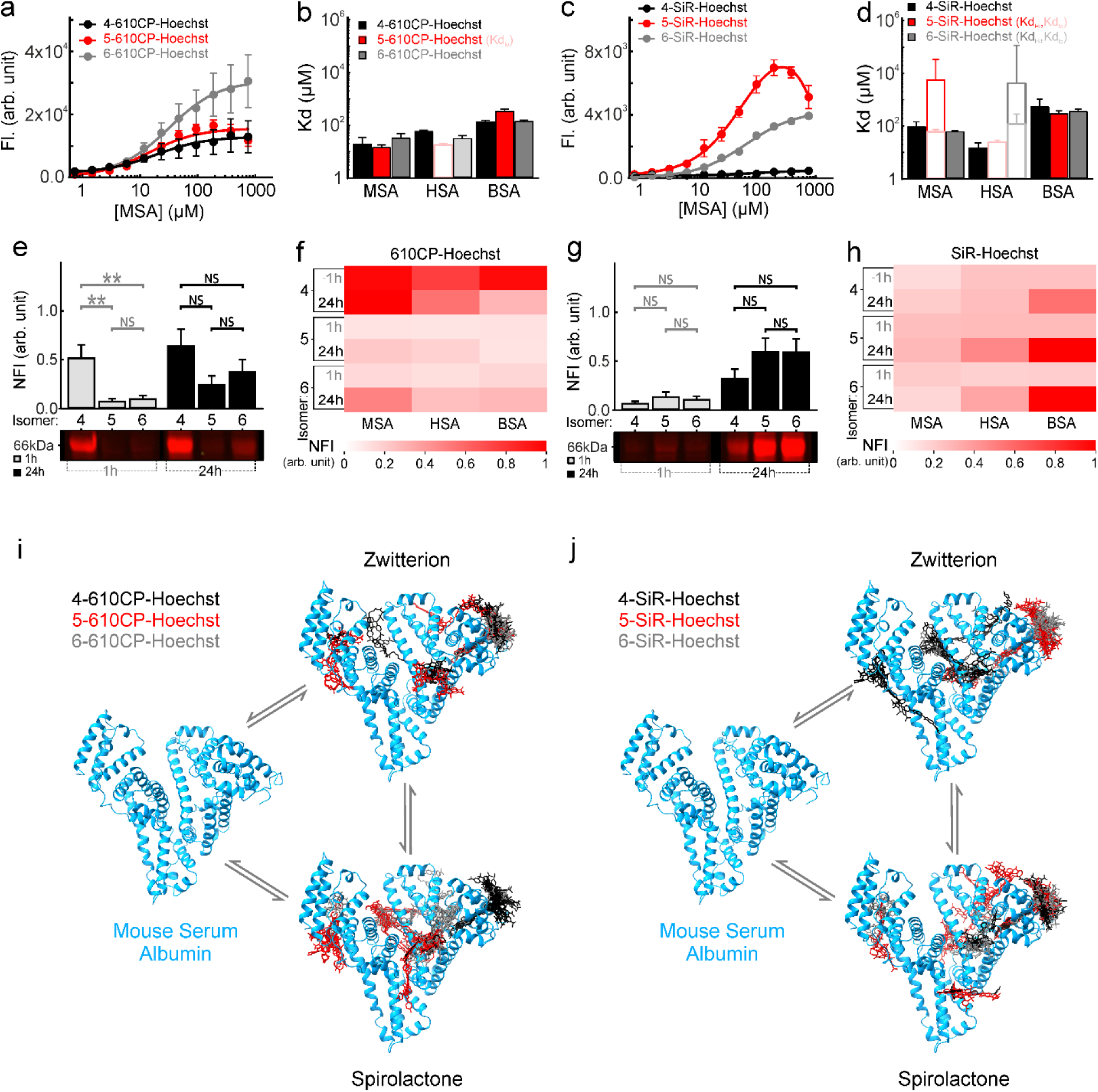
Interaction of Rhodamine-Hoechst probes with serum albumins. Titration of 200 nM probes: 4-, 5-, 6-610CP-Hoechst (**a**) and 4-, 5-, 6-SiR-Hoechst (**c**) with purified mouse serum albumin. Data points of all probes are fitted with a single site-binding equation, except for 5-SiR-Hoechst which was fitted to two site-binding equation. **b, d** Calculated dissociation constants are listed for each regioisomer and three serum albumins (mouse, human and bovine). Data are presented as mean ± s.d of n=3 experimental replicates. **e, g** Covalent binding of 1 µM probes: 4-, 5-, 6-610CP-Hoechst (**e**) and 4-, 5-, 6-SiR-Hoechst (**g**) to serum albumin in mouse plasma. Probes were incubated with plasma purified from mouse blood for 1 h and 24 h, and subsequently subjected to denaturation at 95°C for 5 min followed by SDS-PAGE analysis. The bar graphs indicate the normalized fluorescence intensity of 66 kDa bands on the gels. Data are mean ± s.d of n=5 experimental replicates; ** p≤0.002, NS-not significant in Mann Whitney t-test. **f, h** Heat maps show the average normalized fluorescence intensity of 1 µM DNA probes: 4-, 5-, 6-610CP-Hoechst (**f**) and 4-, 5-, 6-SiR-Hoechst (**h**) covalently bound to purified mouse, human and bovine serum albumin after 1 h and 24 h incubation at 37 °C, and 5 min denaturation at 95 °C. Fluorescence intensities of 66 kDa bands on SDS-PAGE gels were measured and averaged from n=5 independent experiments. MSA, mouse serum albumin; HSA, human serum albumin; BSA, bovine serum albumin; NFI, normalized fluorescence intensity; FS, fluorescence signal. **i**, **j** Molecular docking model of mouse serum albumin bound to all three regioisomers of 610CP- (**i**) and SiR-Hoechst (**j**) existing in an “open” (zwitterion) and “closed” (spirolactone) forms.

Afterwards, we tested the interaction of Hoechst 33342 and free rhodamine dyes with serum albumins (**Figure S5**). The apparent K_d_ values for Hoechst 33342 ranged from 4 to 48 µM (**Tables S3 – S5**) and are similar to previously reported values^[7]^. In contrast to the binding behaviour of Hoechst 33342, we observed a decrease in fluorescence of rhodamine dyes with increasing serum albumin concentration (**Figure S5**). The apparent K_d_ values for 610CP regioisomers were higher (range 10-573 µM) compared to those for SiR dye regioisomers (range 0.16-84 µM) (**Table S6**). These measurements demonstrate that the individual probe components are able to interact with the serum albumin independently.

Next, we measured fluorescence quantum yield and lifetime of the serum albumin-DNA probe complexes. The 610CP-Hoechst and MSA complexes (QY ∼0.05) were significantly brighter compared to the corresponding SiR-Hoechst complexes (QY ∼0.01). Surprisingly, binding to HSA and BSA resulted in the formation of more bright complexes: QY ∼0.2 for 610CP-Hoechst probes and QY ∼0.05 for SiR-Hoechst probes (**Tables S3-S5**). Fluorescence lifetime measurement data could be fitted to two-phase exponential decay. The fast decaying fluorescence (*τ*_1_ ∼0.8 ns) was similar for both fluorophores (610CP and SiR) and it is likely corresponding to intramolecularly quenched state of the probes. The second, slower-decaying fluorescence showed dependence on the rhodamine dye: *τ*_2_∼3.87 ns for 610CP and *τ*_2_ ∼ 3.52 ns for SiR. Interestingly, MSA complexes demonstrated slightly shorter fluorescence lifetimes compared to those with BSA or HSA. The two-phase fluorescence decay was also observed for Hoechst 33342 complexes with all analyzed serum albumins (**Tables S3-S5**). These results are in good agreement with previously reported observations of Hoechst 33258 interaction with BSA ^[7–8]^.

### 2.4. Covalent interaction with serum albumin

After determining how effectively the probes interact with the serum albumins, we next investigated the nature of the interaction between serum albumin and the rhodamine-based Hoechst probes. In this set of experiments, commercially available purified BSA, MSA, and HSA, were incubated with the three regioisomers of 610CP- and SiR-Hoechst for 1 and 24 h at 37 °C in PBS buffer. Following the incubation, samples were denatured at 95 °C and analyzed using SDS-PAGE (**Figure S6**). We assumed that, under such conditions, only covalent complexes will be visible in the in-gel fluorescence images.

First, we performed SDS-PAGE analysis using plasma from mouse blood. It revealed a strong fluorescent band of the 4-610CP-Hoechst-protein crosslink. In contrast, the bands for the 5- and 6’ regioisomers were faint (**Figure 2e**). Quantitative analysis revealed a 10-fold and 6.6-fold increase in fluorescence signal for the 4’ regioisomer compared to the 5’ and 6’ regioisomers, respectively, with no significant difference between the latter two (**Figure 2f**). After 24 hours of incubation, the 4’ regioisomer still displayed a 4.6-fold and 2-fold higher signal compared to the 5’ and 6’ regioisomers (**Figure 2e,f**). Thus, we hypothesized that 4-610CP-Hoechst has a higher propensity to form covalent bonds with MSA. In the case of SiR-Hoechst, no significant differences in fluorescence were detected after 1 hour (**Figure 2g,h**). However, after 24 hours, both 5’ and 6’ regioisomers exhibited a slight fluorescence signal increase of 1.8-fold compared to the 4’ regioisomer (**Figure 2g,h**). We confirmed that tested DNA probes are forming covalent complexes with isolated MSA. The obtained results were identical to data obtained in the isolated plasma (**Figure S6a**,**d**).

In the case of HSA, incubation with 610CP-Hoechst revealed a 7.3-fold and 5.8-fold higher fluorescence for the 4’ regioisomer compared to the 5’ and 6’ regioisomers after 1 hour. At the 24-hour mark, the 4’ regioisomer maintained a 3-fold and 2.2-fold higher signal compared to the 5’ and 6’ regioisomers, respectively (**Figures S6b,e**). For SiR-Hoechst with HSA, after 1 hour, the 5’ regioisomer exhibited a 1.4-fold and 1.5-fold higher signal than the 4’ and 6’ regioisomers, respectively. After 24 hours, this difference increased, with the 5’ regioisomer showing a 3.2-fold and 2.5-fold higher fluorescence compared to the 4’ and 6’ regioisomers (**Figure S6b**,**e**).

Similar trends were observed for BSA. The 4-regioisomer displayed an 8.3-fold and 6.1-fold higher signal compared to the 5- and 6-regioisomers after 1 hour. After 24 hours, both the 4- and 6-regioisomers showed approximately 2.5-fold stronger signals compared to the 5-regioisomer (**Figure S6c**,**f**). For SiR-Hoechst binding with BSA, there was no significant difference in fluorescence at 1 hour, but after 24 hours, the 5-regioisomer showed a slight increase of 1.5-fold increase compared to the 4-regioisomer (**Figure S6c,f**).

These results indicate that 4-610CP-Hoechst probe forms a significant amount of covalent complex with serum albumins. In contrast, 4-SiR-Hoechst is less reactive than the 610CP-based probe, and its covalent binding is close to the background level set by 5’ and 6’ regioisomers. However, we cannot exclude the possibility that covalent complex formation may occur with other proteins present in living cells and organisms.

### 2.5. Modeling of serum albumin and fluorescent probe interaction

The amino acid sequences of albumins of the tested species showed a relatively high 70–75% similarity, however, these minor variations may introduce differences in binding sites and their affinity (**Figure S7**). Thus, we performed molecular docking experiments using X-ray crystallographic data available for BSA (PDB ID: 4JK4) and HSA (PDB ID: 4BKE). MSA crystallography data is not available, however high sequence homology to HSA allowed us to perform homology modeling (template PDB ID: 7OV1) using SWISS-MODEL server^[9]^ and use the obtained structure for the docking experiment. Forty poses of each DNA probe regioisomer in spirolactone and zwitterion form with lowest energy were generated as output for each protein-ligand docking experiment. We identified the most likely binding pockets for each probe, BSA docking results revealed two possible binding sites. DNA probes clustered along central alpha helix (173-204 a.a.) and an additional binding site is composed from several alpha helixes spanning 207-246 a. a. (**Figure S8a**) In HSA docking experiment, 610CP and SiR probes clustered along central alpha helix (174-206 a.a.) (**Figure S8b**). MSA docking experiment yielded three dominating clusters of DNA probe’s binding (**Figure 2i,j**). One of the clusters is located along central alpha helix (198-246 a.a.), the second cluster is surrounded by alpha helixes from 60-100 a.a. and the third cluster is located between two alpha helixes (416-438 and 565-582 a.a.) (**Figure S7a**). This modeling indicates that the performance of the DNA probes, if injected into the blood stream, might be species dependent due to slightly different interaction with serum albumin.

### 2.6. Performance of DNA probes in confocal and STED imaging

Our previous study demonstrated an excellent performance of DNA probes based on Hoechst conjugated to 4- and 5-isomer of rhodamines in human cell lines^[5]^. Herein, we stained mouse NIH3T3 fibroblast with the rhodamine Hoechst probes. We observed that all three regioisomers of 610CP-Hoechst and SiR-Hoechst stained the nuclei of the cells (**Figure S9a,b**). The 5’ regioisomer of 610CP-Hoechst was up to 1.5- and 4-fold brighter comparing to the 4’ and 6’ regioisomer labeled nuclei, as observed in the confocal images. When imaged using STED microscope, a similar pattern was observed with 5’ regioisomer having 5-fold brighter nuclei than 4’ and 6’ regioisomers (**Figure S9c**). In the case of SiR-Hoechst-stained nuclei, the 4’ regioisomer staining of nuclei was brighter than 5’ and 6’ regioisomers for confocal images (**Figure S9a,b**). We also observed several small dotted structures inside the nucleus that were stained distinctly by the three different positional isomers of both 610CP- and SiR-Hoechst. Surprisingly, these structures were the most visible when stained by 4-610CP-Hoechst (**Figure S9a,b,d**). In addition, we tested cytotoxicity of all DNA probes in NIH3T3 fibroblast and found that only relatively high concentrations (>10 µM) of the probes are inducing cell cycle changes (**Figure S10**). These results are similar to the previously reported for human cell lines ^[4a, 5]^.

### 2.7. Performance of DNA probes in two-photon imaging

Next, we hypothesized that the DNA probes could be used for two-photon microscopy which is particularly advantageous for *in vivo* imaging^[10]^. To test this, we performed two-photon microscopy of living human fibroblasts. The 610CP- and SiR-Hoechst were excited at 800 nm and 1300 nm, respectively, using a high power laser. The precise laser power output was measured using a power meter (**Table S7**). Next, we have recorded the laser power vs fluorescence intensity of the stained nuclei (**Figure S11**). The plot for 610CP-Hoechst regioisomers showed that at laser powers below 10 mW, the fluorescence intensity of nuclei increases linearly, but saturation is observed for laser power 10 mW (**Figure S12**). The 6’ regioisomer of 610CP had the highest nucleus fluorescence intensity followed by 4’ and 5’ regioisomers (**Figure S12**). For SiR-Hoechst there is a linear increase in the fluorescent intensity till 5 mW laser power. After that there is a steady increase in the fluorescence intensity till 20 mW laser power (**Figure S12**). At ∼40 mW laser the cells were starting to bleach instantly and the detectors were getting saturated making it impossible to capture the images. Unlike 5’ regioisomer of SiR-Hoechst which gives fluorescence signal at 1 % laser, the 4’ and 6’ regioisomer need 12.5 % and 3 % laser for signal (**Figure S11** and **S12**).

### 2.8. Tail vein injection of fluorescent DNA probes and Imaging of the extracted tissues

Ultimately, we aimed to characterize the performance of the DNA probes in a living organism. We injected the DNA probes into adult CD-1 mice at a dose of 10 mg/kg body weight (**Table S8**). After 24 h, we collected organs: liver, kidney, heart, lungs and brain and analyzed them using microscopy techniques (Figure 3a). Confocal microscopy analysis revealed that 5’ regioisomer of both probes, 610CP- and SiR-Hoechst, selectively stained nuclei in the liver, kidney, heart, lungs (**Figure 3b-d,f-h** and **Figures S13–S19**), while brain tissue was not stained (**Figures S20** and **S21**). In contrast, probes containing 4’ and 6’ regioisomer displayed significant cytosolic staining (**Figures S13–S19**). This observation contrasts with *in cellulo* data, which demonstrate that 4’ regioisomer performs better than 5’ regioisomer ^[11]^.

**Figure 3.**
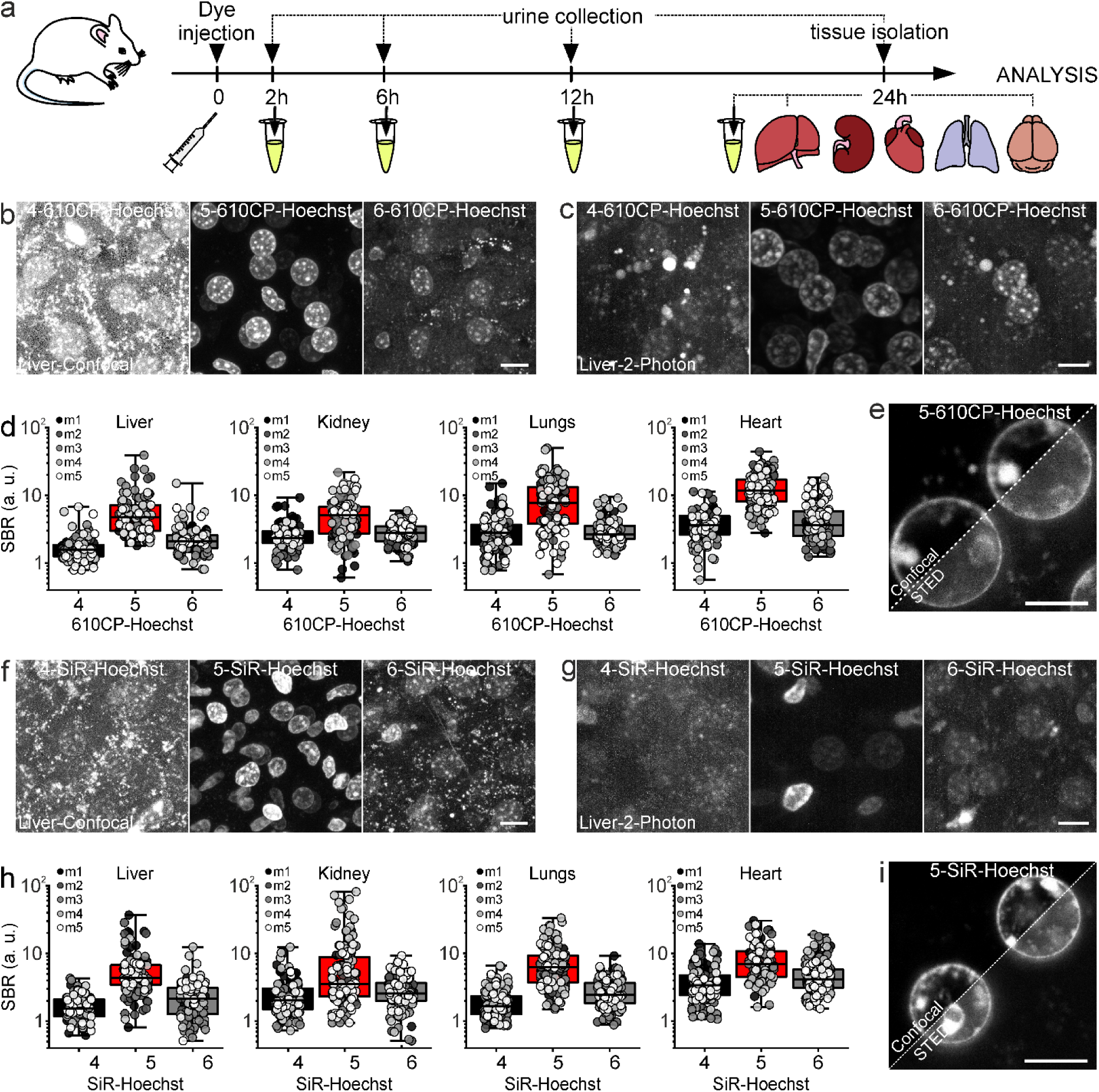
Staining of nuclei by Rhodamine-Hoechst probes in living mice. **a** Graphical representation of methods used to test Rhodamine-Hoechst probes in living mice. CD-1 mice were intravenously administered a single dosage of the probe. To investigate clearance, urine samples were collected at the indicated time points. Living liver, kidney, heart, lungs and brain slices were collected 24 h after injection and immediately subjected to confocal and 2-photon microscopy to investigate the nuclear staining performance. Living liver slices isolated from mice injected with: 4-, 5-, 6-610CP-Hoechst (**b, c**) and 4-, 5-, 6-SiR-Hoechst (**e, f**) and imaged by confocal (**b, f**) and 2-photon (**c, g**) microscope. Scale bar 10 µm. **d, h** Box and strip plots representing signal to background ratio in confocal images of living liver, kidney, lungs and heart slices isolated from mice injected prior 24 h with 4-, 5-, 6-610CP-Hoechst (**d**) and 4-, 5-, 6-SiR-Hoechst (**h**). Data points are signal to background ratios for single nuclei in given organ measured for n=5 animals (measurements for single animal are colour-coded). Averaged signal to background values are indicated by box plot. Numbers 4, 5 and 6 indicate different rhodamine regioisomers. SBR - signal to background ratio. m1-m5 - mouse 1-mouse 5. Confocal and STED imaging of nuclei in liver tissue isolated from mice injected prior 24 h with 5 610CP-Hoechst (**e**) and 5-SiR-Hoechst (**i**). Scale bar 5 µm.

Two-photon excitation microscopy is often used for the visualization of tissues because of its superior penetration depth, better optical sectioning, less light scattering, less background autofluorescence and reduced phototoxicity ^[12]^. Thus, we performed two photon microscopy of tissue samples from liver, kidney, heart, lung, and brain which were isolated from mice injected with different regioisomers of 610CP-Hoechst and SiR-Hoechst DNA probes (**Figure 3a** and **Figures S13 – S19**). The specificity of the nuclear staining was assessed by comparing the staining patterns of different regioisomers across various tissues.

In liver tissue, the 4’ and 6’ regioisomers of 610CP-Hoechst exhibited numerous dotted structures outside the nuclei, indicating nonspecific staining (**Figure 3**). Conversely, the 5’ regioisomer demonstrated specific nuclear staining with minimal background signal, consistent with results obtained from confocal imaging. The robust performance of the 5’ regioisomers facilitated high-resolution imaging of liver nuclei via STED nanoscopy (Figure 3e and l). Similar trends were observed in kidney, heart, and lung tissues (**Figures S13–S19**). The 5’ regioisomer of 610CP-Hoechst and SiR-Hoechst consistently exhibited specific nuclear staining (**Figure 3** and **Figures S13–S19**), while the 4’ and 6’ regioisomers showed nonspecific staining. In heart tissue, nuclei stained with the 4’ regioisomer were not clearly distinguishable, instead appearing as speckled structures, further suggesting nonspecific staining. No efficient staining of nuclei was observed in brain tissues sections with any regioisomer (**Figures S20** and **S21**). In contrast, the strong nuclear DNA signals were observed in brain slices stained after extraction and sectioning (**Figures S22** and **S23**). These results suggest that the blood–brain barrier limits penetration of the DNA probes and that further structural optimization will be necessary to enable effective labeling in the intact brain.

### 2.9. Excretion of fluorescent DNA probes

Fluorescent probes could be excreted via two major pathways–liver and kidney. We have selected to monitor fluorescence in urine of the mice since it allows easy sampling over prolonged periods of time. Thus, we recorded urine fluorescence spectra 2, 6, 12 and 24 h after injection. The results indicate that 24 h after injection most of the fluorescence is clear (**Figure 4** and **Figure S24**). Interestingly, 4-610CP-Hoechst and 4-SiR-Hoechst demonstrated increased excretion after 6 and 12 hours (**Figure 4** and **Figure S24**). The delayed in excretion could be a result of an additional accumulation-retention mechanism which functions in parallel to DNA binding and is consistent with our microscopy imaging findings, which show pronounced off-target staining in addition to nuclear staining. It may also reflect an additional accumulation-retention mechanism, such as the covalent attachment to serum albumin that we have observed.

**Figure 4.**
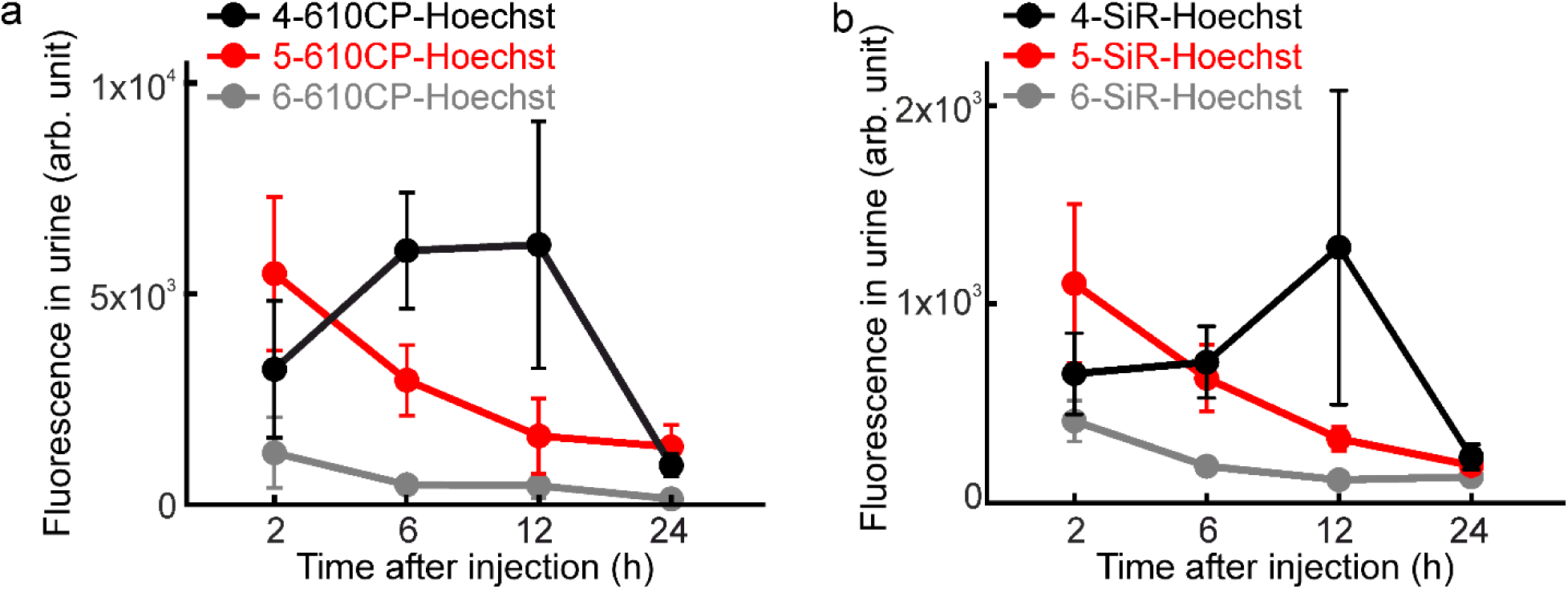
Urine fluorescence after intravenous injection. (**a**) 10 mg/kg of 4-, 5-, 6-610CP-Hoechst or (**b**) 4-, 5-, 6-SiR-Hoechst was injected in the living mouse. Afterwards, urine fluorescence was measured at 2, 6, 12 and 24 h time points in Tecan plate reader. Graphs represent fluorescence at maximum emission wavelength, N ≥ 3, data presented as mean with SEM.

## 3. Conclusion

In this study, we synthesized and evaluated a series of novel red-shifted rhodamine-Hoechst fluorescent DNA probes differing in the positional isomerism of the fluorophore (4-, 5-, and 6-position), with a focus on their photophysical properties, biomolecular interactions, and *in vivo* imaging performance.

The use of Hoechst-C4-NH_2_ as a versatile DNA-targeting moiety enabled efficient coupling to rhodamine dyes via HATU-mediated amide bond formation, yielding the desired conjugates in high yields (60–70%). All three regioisomers (4-, 5-, and 6-position) of the 610CP and SiR derivatives exhibited low QYs in aqueous solution, consistent with our previous observations ^[4a]^. These reductions in fluorescence are attributed to self-aggregation and intramolecular π–π stacking, leading to quenched excited states. This hypothesis is supported by the recovery of fluorescence and absorbance upon SDS addition. Furthermore, the interaction of the DNA probes with hpDNA restored fluorescence intensity and enabled dissociation constant (*K_d_*) determination. Nevertheless, the DNA-bound complexes of the 4’ regioisomers exhibited lower quantum yields compared to their 5’ regioisomers counterparts, potentially reflecting differences in the fluorophore microenvironment. This is further supported by the shorter fluorescence lifetimes observed for both 4-610CP-Hoechst and 4-SiR-Hoechst, which displayed single-exponential decay profiles-suggesting a homogeneous excited state despite reduced brightness.

Binding of the probes to serum albumins (mouse, human, bovine) led to fluorescence activation, most likely via spirolactone ring opening and relief of intramolecular quenching. Binding affinity analysis revealed that 610CP and SiR regioisomers generally fit a single-site model (4′ and 6′ regioisomers), while the 5’ regioisomers exhibited dual binding modes, with a secondary low-affinity interaction that contributed to quenching. Quantum yield and lifetime data indicated that the brightness of albumin-bound 610CP probes (QY ∼0.2 for HSA/BSA complexes) exceeded that of the SiR derivatives (QY ∼0.05). Two-phase exponential fluorescence decay was observed for all albumin-probe complexes, consistent with a mixture of quenched and unquenched probe populations. SDS-PAGE analysis revealed distinct differences in covalent crosslinking efficiency among the regioisomers. The 4-610CP-Hoechst displayed a markedly higher propensity for covalent binding to MSA, BSA, and HSA than the 5’ and 6’ regioisomers, suggesting enhanced electrophilicity or proximity-induced reactivity at the 4-position. We propose that the covalent dye–protein crosslinking arises from the formation of a reactive anhydride intermediate, a mechanism previously reported for N-alkyl-substituted phthalamic acids^[13]^. In contrast, 4-SiR-Hoechst showed minimal covalent labeling, indicating probe-dependent reactivity likely influenced by the electronic properties of the dye scaffold. All of these observations were supported by molecular docking experiments which revealed species-specific differences in binding site architecture and albumin variant-dependent clustering of binding sites for the positional isomers.

The 5-610CP-Hoechst consistently yielded the brightest nuclear staining of mouse NIH3T3 fibroblasts in both confocal and STED modalities. Conversely, 4-SiR-Hoechst displayed slightly superior brightness compared to 5-SiR-Hoechst and 6-SiR-Hoechst. Intriguingly, 4-610CP-Hoechst highlighted intranuclear punctate structures, suggesting unique binding or accumulation properties not observed with other regioisomers. In two-photon microscopy, fluorescence response to increasing laser power revealed a different picture. 6-610CP-Hoechst displayed the brightest signal at low laser powers, suggesting favorable properties, while 5-SiR-Hoechst was the brightest among the SiR regioisomers.

Tail vein injection into mice revealed rhodamine regioisomer-specific differences in probe localization. The 5’ regioisomers of both 610CP- and SiR-Hoechst selectively stained nuclei in multiple tissues (liver, kidney, lung, heart), with minimal cytosolic background and no signal in the brain, suggesting blood-brain barrier impermeability. In contrast, 4’ and 6’ regioisomers exhibited substantial cytosolic staining, implying nonspecific retention or off-target interactions. These discrepancies between *in vitro* and *in vivo* nuclear specificity underscore the importance of protein binding and clearance mechanisms in probe biodistribution. Two-photon imaging of tissue slices confirmed that the 5’ regioisomers maintain nuclear specificity in various organs, whereas the 4’ and 6’ regioisomers displayed significant nonspecific background. The absence of staining in brain tissue highlights a critical limitation for central nervous system imaging applications and calls for structural modification to enhance brain-blood barrier penetration.

Urine fluorescence monitoring revealed that the 4’ regioisomers of both 610CP and SiR were rapidly excreted after 6-12 hours post-injection. It might be consequence of the observed off-target staining and possible covalent interaction with serum albumin. In contrast, the 5’ regioisomers demonstrate superior nuclear specificity, show no delay in excretion, and benefit from the rapid clearance of excess dye. These findings illustrate the trade-off between probe retention off-targeting and high specificity staining *in vivo*.

In conclusion, this comprehensive evaluation reveals that the positional isomerism of rhodamine conjugation to Hoechst strongly influences the photophysical behavior, protein binding profile, imaging performance, and *in vivo* biodistribution of DNA-targeted fluorescent probes. The 5’ regioisomer consistently emerges as the most suitable candidate for high-specificity nuclear imaging in cells and tissues, demonstrating optimal performance in both confocal and super-resolution modalities. Meanwhile, the 4’ regioisomer exhibits enhanced covalent binding capacity but suffers from higher nonspecific background and delayed systemic clearance. Although these properties could, in principle, lead to adverse effects such as immune responses, such reactions are unlikely to occur during acute staining experiments. Our results underscore the critical role of fluorophore orientation and the local environment in determining the behaviour of DNA probes and emphasizing the need for careful molecular design tailored to specific imaging applications. Future efforts should focus on engineering rhodamine derivatives with optimized brightness, improved permeability and enhanced *in vivo* stability to expand the utility of these probes for real-time DNA imaging in complex biological systems.

## 4. Experimental section

### Materials

All chemicals were purchased from Sigma-Aldrich (Munich, Germany), Merck (Darmstadt, Germany) or TCI (Eschborn, Germany) and used as received. Deuterated solvents for NMR spectra recording were purchased from Deutero GmbH (Kastellaun, Germany). Probes were dissolved in anhydrous DMSO (Sigma-Aldrich cat. 900645-4X2ML) as 1000× stocks and stored at -20°C. The following purified serum albumins were used: bovine serum albumin (Sigma-Aldrich, cat. A7030), human serum albumin (Sigma-Aldrich, cat. A1653) and mouse serum albumin (Loxo, cat. IMSALB1000MG).

### Preparation of hairpin DNA

The hairpin forming oligonucleotide 5’-CGCGAATTCGCGTTTTCGCGAATTCGCG-3’ (28 nt) for DNA binding studies was purchased from Sigma-Aldrich^[4b]^. The lyophilized oligonucleotide was dissolved in PBS (Lonza, Cat. No. BE17-516F) at 1 mM concentration (hpDNA solution). Hairpin was formed by putting the tube with hpDNA solution into a boiling water bath and subsequently slowly cooling it down to room temperature (RT).

### Plasma isolation

0.5 M EDTA (pH 8.0) treated mouse blood was centrifuged at 4 °C and 1200 x g for 10 min. Clear, plasma containing supernatant was harvested and stored at -20 °C until further use.

### Absorbance and fluorescence spectra measurements

Absorbance and fluorescence spectra measurements of 2 μM Rhodamine-Hoechst probes in PBS (Lonza, Cat. No. BE17-516F), PBS containing 30 μM hpDNA, PBS containing 6.25mg/ml serum albumin and PBS containing 0.1% SDS were performed. 2 μM probes were incubated at RT for 2 h before measurements. The mixtures were incubated at RT for 2 h before measurements.

The solutions were pipetted to a glass bottom 96-well plates (MatTek, Cat. PBK96G-1.5-5-F). Absorbance and fluorescence were measured using a Tecan Spark® 20M multimode microplate reader. The absorbance was recorded at 320-850 nm. Solutions with 610CP-Hoechst were excited at 565 nm and the ones with SiR-Hoechst at 610 nm, while emission was recorded at 600-850 nm and 620-850 nm, respectively, with a step size of 2 nm. The excitation and emission bandwidths were 10 nm and 5 nm for both groups of probes. Gain for 610CP-Hoechst and SiR-Hoechst was set to 100 and 110 respectively, to ensure optimal signal detection. The Z-position was set to 18900 µm with an integration time of 40 µs. Interpolation of result was performed using graph pad prism software (version 10.3.0), and the results were expressed as mean ± SD. Experiments were performed in triplicates (N=3).

### Determination of quantum yields and lifetimes

The fluorescence absolute quantum yields (QY) values were obtained with a Quantaurus-QY spectrometer (model C11347-12, Hamamatsu) according to the manufacturer’s instructions. For solutions containing 610CP-Hoechst, the excitation wavelength was set to 570 nm, and for solutions containing SiR-Hoechst probes, it was set to 620 nm. Emission was recorded at 600-850 nm and 654-860 nm for 610CP-Hoechst and SiR-Hoechst, respectively.

Fluorescence lifetimes were measured with a Quantaurus-Tau fluorescence lifetime spectrometer (model C11367-32, Hamamatsu) according to the manufacturer’s instructions. For 610CP-Hoechst excitation and emission wavelengths were set at 590 nm and 640 nm, respectively. For SiR-Hoechst excitation and emission wavelengths were set at 630 nm and 670 nm, respectively. The time range was set as 53 ns. For the spectroscopy measurements of probe-hpDNA interaction (5’ and 6’ regioisomers of 610CP- and Sir-Hoechst) all solutions were performed in air-saturated PBS buffer containing 2 μM probe and 30 μM hairpin-forming oligonucleotide after incubation for 2 h at RT. For 4’ regioisomers of rhodamine probes, 1 μM probe and 20 μM hairpin-forming oligonucleotide was used. Similarly, for probe-serum albumin interaction, the solutions containing 1 uM probe with 25 mg/ml Serum Albumin in PBS buffer were used, which were incubated for 1 h at RT before measurement. Analysis of the results was performed using graph pad prism software (version 10.3.0), and the data was presented as mean ± SEM, N = 3.

### Determination of Kds

Hoechst 33342, 610CP- and SiR probes or dyes were prepared in PBS (Lonza, cat. BE17-516F) and dispensed into glass-bottom 96-well plates (MatTek, cat. PBK96G-1.5-5-F). For DNA binding assays, probes were titrated with increasing concentrations of hpDNA. For serum albumin binding assays, probes were titrated with increasing concentrations of mouse serum albumin, human serum albumin, or bovine serum albumin in PBS. The plates were incubated for 1 h at RT prior to fluorescence measurement on a Tecan Spark® 20M multimode microplate reader. Hoechst 33342, 610CP- and SiR-fluorophores were excited at 360 nm, 570 nm and 640 nm while emission was recorded at 380-650 nm, 600-850 nm and 640-850 nm, respectively, with a step size of 2 nm. The excitation and emission bandwidths were 10 nm and 5 nm for all. Gain was set to 120 for Hoechst and 160 for 610CP Hoechst and SiR Hoechst and the Z – position was set to 18900 µm with an integration time of 40 µs. To study dye-serum albumin and probe-serum albumin binding interaction, mouse, human and bovine serum albumin concentrations were varied from 0 µM to 751.8 µM for a fixed concentration of dyes (2 µM) and probes (200 nM). The apparent K_d_ values were determined as previously described by plotting the fluorescence intensity of the emission signal versus hpDNA or serum albumin concentration ^[4a]^. The data points were fitted using GraphPad Prism (version 10.3.0) to the “Two site-binding + offset” or “Single site binding” function. Samples were measured in triplicates (N=3) and the experiment was repeated on different days. The data are represented as mean ± SD, fitted Kd values presented as mean ± SEM.

### Crosslinking of the probes to serum albumin

A solution of mouse plasma, mouse, human and bovine serum albumin (1.56 mg/ml) in PBS (Lonza, cat. BE17-516F) was incubated with the DNA probes (2 µM) or DMSO for 1 h and 24h at 37 °C. Afterwards, 4x sample buffer (0.2 M Tris-HCl (pH 6.8), 8 % SDS (w/v), 25 % glycerol (v/v), 6 mM Bromophenol Blue and 5% 2-mercapto-ethanol) was added to the incubated solution and the sample was heated at 95°C for 5 min. Then, the sample was loaded onto BioRad Mini Protean TGX gel (4-15 %; BioRad, cat. 456-1086) and separated in running buffer (0.25 M Tris, 1.925 M glycine, 1 % SDS pH 8.3) under 150 V for 1 h. After that, fluorescence signal from the gel was detected by Amersham Imager 600 RGB. Subsequently, the gel was incubated in Coomassie stain (20 % 5 x Roti-Blue ((v, v); Roth, cat. A152.1), 20 % MeOH in H_2_O) for 2 h, ensuring thorough coverage, de-stained in solution (10 % acetic acid (v, v) in H_2_O) for 3 h and imaged again thereafter. All measurements were performed 5 times on different days (N=5), the data analysed using GraphPad Prism software (version 10.3.0) and the results are shown as mean ± SD. The statistical analysis was performed through Mann Whitney t-test and P ≤ 0.05 and P ≤ 0.002 was considered significant.

### Molecular Docking

For docking, X-ray structures of Human serum albumin PDB ID 4BKE^[14]^ and Bovine serum albumin PDB ID 4JK4^[15]^ were used. Mouse serum albumin complex structure modelling was performed using Swiss-Model server. The DNA-probes were drawn using ChemDraw Professional 15.1 and prepared for docking with AutoDock Tools version 1.5.6 ^[16]^. The docking simulation was performed using Vina Autodock version 1.2.0 ^[17]^. Amino acids interacting with ligand were identified using LigPlot+ version 2.3 ^[18]^.

### Maintenance of cultured cells

NIH3T3 mouse fibroblasts (ATCC-CRL-1658) were cultured in High Glucose DMEM (Thermo Fisher Scientific, cat. 31966047) supplemented with 10 % (v/v) FBS (Bio&Sell, cat. S0615) and 1 % (v/v) Penicillin-Streptomycin (Sigma, cat. P0781) in a humidified 5 % CO_2_ incubator at 37 °C. Cells were seeded on coverslips in polystyrol culture plates 24 h before further experimentation.

Human fibroblasts were cultured in high-glucose DMEM (Thermo Fisher, #31053044) and 10% FBS (Thermo Fisher, #10082147), 1% sodium pyruvate (Sigma, #S8636), 1% GlutaMax (Thermo Fisher, # 35050038) in the presence of 1% penicillin–streptomycin (Sigma, #P0781) in a humidified 5% CO2 incubator at 37 °C. Cells were seeded in glass bottom 6-well plates (MatTek, #P06G-1.0-20-F) 24h before imaging experiments.

### Maintenance of animals

Animal procedures were carried out in accordance with institutional regulations on animal use in research. Experiments performed on living animals were approved and authorized by the Lower Saxony State Office for Consumer Protection and Food Safety (Niedersächsisches Landesamt für Verbraucherschutz und Lebensmittelsicherheit). Sacrificing rodents for subsequent organ isolation followed by preparation of living slices and cultures did not require specific authorization (Animal Welfare Law of the Federal Republic of Germany, Tierschutzgesetz der Bundesrepublik Deutschland (TierSchG); §7 Abs. 2 Satz 3 TierSchG). The animals used for the experimentation were CD-1 wild type male and female mice more than 8 weeks old and weighing about 35-45 grams. All mice were housed with a 12 hours light/dark cycle and free access to food and water.

### Cytotoxicity study by two-step cell cycle analysis

Protocol of cytotoxicity study using NucleoCounter® NC-3000™ two-step cell cycle analysis has been described previously ^[4a]^. Briefly, serial dilutions of the probes (concentration: 20, 10, 5, 2.5, 1.25, 0.625, 0.3125 and 0 µM; stock solutions of probes were prepared in DMSO) were added into the media of cultured NIH3T3 cells (6-well plates, ∼250,000 cells per well) and incubated for 24 h at 37 °C in a humidified incubator with 5% CO_2_. Then, the cells were incubated at 37 °C for 5 min with 250 μl lysis solution (Solution 10, Chemometec Cat. No. 910-3010) supplemented with 10 μg/ml DAPI (Solution 12, Chemometec Cat. No. 910-3012) per well. After that, 250 μl of stabilization solution (Solution 11, Chemometec Cat. No. 910-3011) was added. Next, the chamber of a NC-Slide A2™ (Chemometec, Cat. No. 942-0001) was loaded with ∼30μL of the cell suspension and the DNA content of ∼10000 cells and measured using the NucleoCounter® NC-3000™ System. Acquired cell cycle histograms were analyzed with ChemoMetec NucleoView NC-3000 software, version 2.1.25.8. The experiments were repeated five times on different days. The results presented are mean ± s. d..

### Staining of nuclei in living cells

Cells were stained in ibidi µ-Slide 8 Well ^high^ Glass Bottom plate (IBIDI, # 80807) with the fluorescent probes in DMEM (Thermo Fisher, #31053044) supplemented with 10% FBS (Bio&SELL, #S0615) for 1h at 37 °C and 5% CO_2_. The cells were washed 2 times with media and imaged in DMEM with 10 % FBS. The nuclei of the living cells were imaged with confocal, STED or two-photon microscope. No-wash experiments were performed the growth medium after probe addition and incubation for the indicated period of time. At least 12 nuclei (N≥12) were acquired and considered for data analysis. The image was processed using FIJI (Image J version 1.54g) and the data obtained was analyzed with graph pad prism software (version 10.3.0). The results were expressed as mean ± SEM. The statistical analysis was performed through Mann Whitney t-test and P < 0.05, 0.002, 0.0002 and 0.0001 was considered significant.

### Preparation of living tissue slices

Injected mice were sedated with isoflurane in a sealed container and quickly euthanized by cervical dislocation. The skull and the abdomen were opened. The organs: liver, kidney, heart, lungs and brain were taken out instantly, transferred to ice cold EGTA solution ^[19]^ and sliced manually using a scalpel blade under a binocular microscope. Prepared tissue slices were then put on a slide with EGTA solution and coverslip was gently mounted on the slide and sealed. The tissues were imaged immediately after by Leica TCS SP8 confocal, LaVision TriM Scope II Multiphoton microscope or Abberior Expert Line nanoscope. One lobe of liver was snap frozen in liquid nitrogen and stored at -80°C for tissue proteome analysis.

For *ex vivo* staining of the brain slices, adult CD-1 mice of both genders were sedated with isoflurane in a sealed container and quickly euthanized by cervical dislocation. Brains were dissected instantly, transferred into ice cold EGTA (pH 7.4) infused with 95 % O2 and 5 % CO2 and sliced into 300 μm thick coronal sections using a vibratome (Leica VT1200S) straight away. Prepared slices were incubated in EGTA solution containing 250 nM probes on ice for 30 min. and imaged by Leica TCS SP8 confocal microscope or Abberior Expert Line nanoscope without washing at RT thereafter.

### Injection of the fluorescent probes

Adult CD-1 mice of both genders were carefully fixed in a restrainer (Table S8), and injected intravenously (i.v.) with the rhodamine probes in DMSO (10 mg/kg body weight) along with saline (sterile 0.9% NaCl) and Pluronic F-127 (Table S9). Afterwards, animals were placed back in the cage and allowed to recover from the injection. The optimum dose for probes to be injected was selected from the previous literature ^[20]^.

### Confocal microscopy

Confocal images of liver, kidney, heart, lungs and brain were acquired with a TCS SP8 (Leica) microscope with a HC PL APO 20x/0.70 IMM (Leica) as well as a HC PL APO CS2 63x/1.40 Oil (Leica) objectives. Fluorophores were excited using 561 nm and 633 nm excitation lasers. Photomultiplier tube detectors were used to detect the fluorescence. Image acquisition parameters are listed in **Table S10**.

### STED microscopy

STED nanoscopy was performed using ExpertLine STED microscope (Abberior Instruments GmbH) equipped with an UPlanSApo 100x/1.40 oil objective (Olympus), 561 nm and 640 nm 40 MHz pulsed excitation laser lines and a pulsed 775 nm 40 MHz 3 W depletion laser. Detection bandwidth of the APD filters was 650-720 nm. The pinhole was set to 0.9 Airy units. A pixel size of 20 nm was set with a pixel dwell time of 15 µsec and a line accumulation of 2. The STED power was set to 30 %. The excitation power was set depending on the sample brightness and kept constant per experiment.

### Image processing

All acquired images were processed and visualized using Fiji ^[21]^. LIF files of the confocal images and MSR files for the images acquired on STED microscopy were imported using bio-formats importer plugin to Fiji. In the confocal images nuclei was marked as the region of interest and the fluorescence intensity was measured. For measuring the background ‘make inverse’ tool was used and the region outside the nuclei was measured. The data acquired was analyzed in GraphPad Prism (version 10.3.0). The signal to background ratio was plotted using standard python libraries (matplotlib, seaborn). All measurements acquired on confocal microscope were recorded from 5 mice and at least 100 nuclei (N ≥ 100) were considered for analysis. The data was represented as mean ± SD. The statistical analysis was performed through Mann Whitney t-test and P < 0.05 (*), 0.002 (**), 0.0002 (***) and 0.0001 (****) was considered significant.

### Two Photon power measurement

For laser power measurement the ‘blanking’ of the scan mirrors in the scan head on the microscope was disabled assuring a fixed/‘parked’ position of the laser. The laser shutters in the system were opened and the laser beam in the objective plane (Olympus 25x/ 1.05 XLPLN25XWMP2, NA:1.05, water #N5165100) was measured using a laser power sensor from Ophir (3A-PF-12) with laser power meter Ophir Nova II (Nova II monochrome LCD laser power and energy meter). The laser power was measured at 1, 3, 6.5, 12.5, 25, 50 and 100% laser and wavelength 800 and 1300nm (**Table S7**).

### Two-photon microscopy of human fibroblasts

Human fibroblast cells incubated with 200 nM rhodamine probes for 1 hr at 37°C. The cells were washed with media before being imaged using two-photon microscope. The measurements were performed at 1, 3, 6.5, 12.5, 25 and 50 % laser with a nominal power of ∼0.891W for 800 nm and ∼0.915W for 1300 nm. Images were acquired using an Olympus 25X (NA 1.05, water #N5165100) objective. The backscattered emitted light was split by 495 nm and 560 nm long pass dichroic mirrors (Semrock) and collected by photomultiplier tube (PMT) detectors (H6780-01-LV, Hamamatsu, Herrsching am Ammersee, Germany). Signals for cells incubated with 610CP-Hoechst and SiR-Hoechst were imaged using the filter settings 620 ± 60 nm and 651LP (BrightLine HD filters, Semrock, AHF Analysentechnik Tübingen, Germany), respectively. Snapshot measurements were acquired with Individual image sizes of 112.3 × 112.3 µm with 1024 × 1024 pixels.

### Two-photon microscopy of tissues

Two-photon excited fluorescence (2PEF) images were acquired using an upright multiphoton microscope TriM Scope II (Miltenyi Biotec, Bielefeld, Germany) equipped with a tunable femtosecond laser and two independently tunable output channels (Cronus 2P, Light Conversion, Vilnius, Lithuania). The first (680–960 nm) and second tunable channel (950–1300 nm) were used for exciting tissue samples injected with 610CP-Hoechst and SiR-Hoechst, respectively.

610CP-Hoechst and SiR-Hoechst were excited at 800 nm and 1300 nm, respectively. The laser operated between 1-80 % of its maximal output power (∼0.891W for 800 nm and ∼0.915W for 1300 nm; **Table S11**). Images were acquired using a Zeiss W Plan-Apochromat 20× (NA 1.0) water immersion objective and Olympus 25X (NA 1.05, water #N5165100). The backscattered emitted light was split by 495 nm and 560 nm long pass dichroic mirrors (Semrock) and collected by photomultiplier tube (PMT) detectors (H6780-01-LV, Hamamatsu, Herrsching am Ammersee, Germany). Signals for tissue samples injected with 610CP-Hoechst and SiR-Hoechst were imaged using the filter settings 620±60 nm and 647LP (check manufacturer) (BrightLine HD filters, Semrock, AHF Analysentechnik Tübingen, Germany), respectively. Z stacks of sections were acquired with individual image sizes of 56×56 µm with 1024×1024 pixels, a pixel dwell time of 0.64 µs.

## Acknowledgement

The authors thank the Max Planck Society for supporting this work. The authors thank Prof. Stefan W. Hell for his support. SP was supported by the International Max Planck Research School for Genome Science, part of the Göttingen Graduate School for Neurosciences, Biophysics, and Molecular Biosciences. The authors thank the animal house facility at MPI-NAT Fassberg-Campus for their support with animal experiments. The authors thank Dr. Mišo Mitkovski and Heiko Röhse (Light Microscopy Facility at City-Campus, MPI-NAT) for their assistance with two-photon microscopy imaging. The authors are grateful to Dr. Vladimir Belov, Jan Seikowski, Jens Schimpfhauser and Jürgen Bienert (Facility for Synthetic Chemistry, MPI-NAT) for the NMR measurements and the central analytics’ team (Institute for Organic and Biomolecular Chemistry, Georg-August University, Göttingen) for acquiring HRMS. The authors are grateful to Dr. Julien Guy for his support with Python and data visualization. Figures 2i,j and S8 were created using UCSF ChimeraX, developed by the Resource for Biocomputing, Visualization, and Informatics at the University of California, San Francisco, with support from National Institutes of Health R01-GM129325 and the Office of Cyber Infrastructure and Computational Biology, National Institute of Allergy and Infectious Diseases.

## Author contributions

S.P., K.A.K. and G.L. conceived and planned the study. S.P, K.A.K., J.B., T.K. and G.L. performed the experiments. S.P, K.A.K., J.B., and G.L. performed the data analysis. S.P, K.A.K. and G.L. wrote the initial draft; all authors contributed to the final version of the manuscript.

## Data availability

The data that support the findings of this study are available from the corresponding author upon reasonable request.

