## Supplementary information for "Regioisomer-controlled red-shifted DNA probes for imaging of living tissues"

#### General chemical experimental information

NMR spectra were recorded at 25 °C with an Agilent 400-MR spectrometer at 400.06 MHz ($^1$H) and 100.60 MHz ($^{13}$C) and are reported in ppm. All $^1$H and $^{13}$C spectra are referenced to tetramethylsilane (δ = 0 ppm) using the residual signals of the solvents according to the values reported in literature[1]. Multiplicities of signals are described as follows: s = singlet, d = doublet, t = triplet, q = quartet, p = pentet, m = multiplet or overlap of non-equivalent resonances, br = broad signal. Coupling constants (J) are given in Hz. All NMR spectra were processed with MestRenova 11.0.4 software.

ESI-HRMS were recorded on a MICROTOF spectrometer (Bruker) equipped with ESI ion source (Apollo) and direct injector with LC autosampler Agilent RR 1200. The mass scanning range was set as 50-1600 m/z and the dry gas temperature was 180 °C. The capillary voltage was set to 4.5 kV. The dry gas was set to 4.0 l/min.

Analytical LC–MS analysis was performed on an Agilent 1260 Infinity II LC/MS system equipped with an autosampler, diode array detector WR, fluorescence detector Spectra and Infinity Lab LC/MSD 6100 series quadruple with API electrospray with a mass resolution of 0.1 Da. Analysis was done by using an Ascentis Express 90 Å AQ-C18, 5 cm x 2.1 mm, 2 µm column with A: 25 mM $HCOONH_4$ (pH = 3.6) aqueous buffer and B: MeOH. Only positive ion mode was used for sample analysis. The mass scanning range was 100-2000 m/z and the dry gas temperature was 350 °C. The voltage was set to 4.5 kV. Absorption was monitored at 254 nm, 650nm (for 4-SiR-Hoechst), and 254nm, 610 nm (for 4-610CP-Hoechst).

Preparative HPLC was performed on a combined Agilent 1290 Infinity II preparative system equipped with a 1290 Infinity II open-bed sampler (G7169B)/fraction collector (G7159B), 1260 Infinity II preparative binary pump (G7161A), 1260 Infinity II multiple wavelength detector (G7165A) and with Agilent Pursuit 10 C18, 10 µm, 250 x 50 mm preparative column.

**Synthesis of the probes**

For the preparation of 4-SiR/4-610CP-Hoechst into a solution of corresponding rhodamine dye (30 µmol, 1 eq; 4-SiR-COOH [2] or 4-610CP-COOH [2] and DIPEA (25 µL) in DMSO (300 µL) a solution of HATU (39 µmol, 1.3 eq) in DMSO (200 µL) was added and the mixture was mixed for 3 min. Afterwards a solution of Hoechst-$C_4$-$NH_2$ [3] (45 µmol ,1.5 eq) in DMSO (250 µL + 15 µL DIPEA) was added at once. Reactions were stirred for 30 min. and their course was monitored by LC/MS analysis. Once reaction was finished it was quenched with 40 µL of formic acid, diluted with water and acetonitrile to 3 mL volume and further purified by the means of preparative HPLC (preparative column: Agilent Pursuit 10 C18, 10 µm, 250 x 50 mm).

4-610CP-Hoechst:

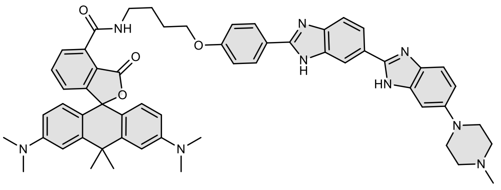

Compound was purified by preparative HPLC (solvent A: $H_2O$ + 0.2 % HCOOH, solvent B: MeCN; temperature 25 °C, gradient A:B - 3 min. 70:30 isocratic, 4-20 min. 70:30 to 0:100 gradient, 20-25 min. 0:100 isocratic, 150 mL/min. flow). Fractions containing the product were collected, evaporated and lyophilized from acetonitrile water mixture. 16.5 mg of blue solid was obtained with a yield of 59%.

$^1$H NMR (400 MHz, Methanol-$d_4$ + $CF_3COOD$) δ 8.48 (d, *J* = 1.7 Hz, 1H), 8.19 – 8.10 (m, 3H), 8.08 (d, *J* = 5.1 Hz, 1H), 7.98 (d, *J* = 8.6 Hz, 1H), 7.82 – 7.72 (m, 3H), 7.46 – 7.38 (m, 2H), 7.34 (d, *J* = 2.2 Hz, 1H), 7.25 (d, *J* = 2.6 Hz, 2H), 7.24 – 7.20 (m, 2H), 7.11 (d, *J* = 9.4 Hz, 2H), 6.84 (dd, *J* = 9.4, 2.6 Hz, 2H), 4.18 (t, *J* = 6.1 Hz, 2H), 3.97 (d, *J* = 13.2 Hz, 2H), 3.69 (d, *J* = 12.1 Hz, 2H), 3.50 (t, *J* = 6.8 Hz, 2H), 3.35 (d, *J* = 6.8 Hz, 2H), 3.23 (q, *J* = 11.7, 10.5 Hz, 2H), 3.01 (s, 3H), 2.01 – 1.93 (m, 2H), 1.91 – 1.83 (m, 5H), 1.74 (s, 3H) (Fig. S2).

$^{13}$C NMR (101 MHz, Methanol-$d_4$ + $CF_3COOD$) δ 170.93, 169.56, 164.71, 163.28, 158.30, 157.60, 157.30, 155.13, 150.82, 150.11, 138.88, 138.69, 137.94, 136.82, 134.71, 132.25, 131.74, 130.94, 129.58, 128.22, 125.09, 122.12, 120.65, 119.64, 117.90, 116.81, 116.76, 116.47, 115.65, 115.23, 114.07, 112.24, 101.05, 69.30, 68.88, 54.62, 48.49, 43.57, 42.92, 41.14, 41.03, 40.69, 35.95, 32.50, 27.59, 26.91 (Fig S2).

ESI-MS, positive mode: m/z = 934.5 $[M+H]^+$. HRMS (ESI) calculated for $C_{57}H_{60}N_9O_4$ $[M+H]^+$ 934.4763, found 934.4763.

4-SiR-Hoechst:

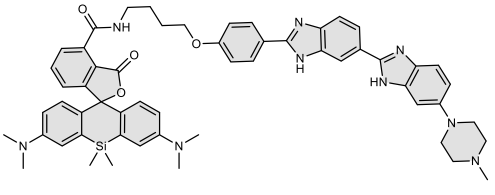

Compound was purified by preparative HPLC (solvent A: $H_2O$ + 0.2 % HCOOH, solvent B: MeCN; temperature 25 °C, gradient A:B - 3 min. 40:60 isocratic, 4-20 min. 40:60 to 0:100 gradient, 20-25 min. 0:100 isocratic, 150 mL/min. flow). Fractions containing the product were collected, evaporated and lyophilized from acetonitrile water mixture. 20 mg of light blue solid was obtained with a yield of 70%.

$^1$H NMR (400 MHz, DMSO-$d_6$ + $CF_3COOD$) δ 9.1 (t, *J* = 5.6 Hz, 1H), 8.3 (s, 1H), 8.1 (d, *J* = 8.8 Hz, 2H), 8.0 (d, *J* = 8.6 Hz, 1H), 7.8 – 7.7 (m, 2H), 7.7 (d, *J* = 8.4 Hz, 1H), 7.4 (d, *J* = 8.7 Hz, 1H), 7.3 – 7.2 (m, 1H), 7.2 – 7.1 (m, 2H), 7.0 (d, *J* = 2.8 Hz, 3H), 6.9 (dd, *J* = 8.8, 2.2 Hz, 1H), 6.7 (d, *J* = 8.9 Hz, 2H), 6.6 (dd, *J* = 9.1, 2.8 Hz, 2H), 4.1 (t, *J* = 6.3 Hz, 2H), 3.4 (q, *J* = 5.9 Hz, 4H), 3.1 (t, *J* = 4.9 Hz, 4H), 2.9 (s, 12H), 2.3 (s, 3H), 1.9 (p, *J* = 6.5 Hz, 2H), 1.8 (p, *J* = 7.1 Hz, 2H), 0.6 (s, 3H), 0.5 (s, 3H) (Fig. S1).

$^{13}$C NMR (101 MHz, DMSO + + $CF_3COOD$) δ 169.62, 169.09, 165.22, 165.18, 164.13, 164.09, 164.07, 160.72, 157.71, 157.02, 156.07, 153.26, 149.64, 148.16, 148.12, 136.26, 136.10, 135.01, 131.01, 129.33, 128.71, 128.68, 128.40, 125.80, 124.85, 122.72, 121.97, 121.86, 116.71, 115.35, 114.17, 114.10, 90.71, 67.97, 55.34, 50.41, 46.22, 40.26, 26.69, 26.04, 0.46, -0.68 (Fig. S1).

ESI-MS, positive mode: m/z = 950.5 $[M+H]^+$. HRMS (ESI) calculated for $C_{56}H_{60}N_9O_4Si$ $[M+H]^+$ 950.4532, found 950.4532.

#### LC-MS data

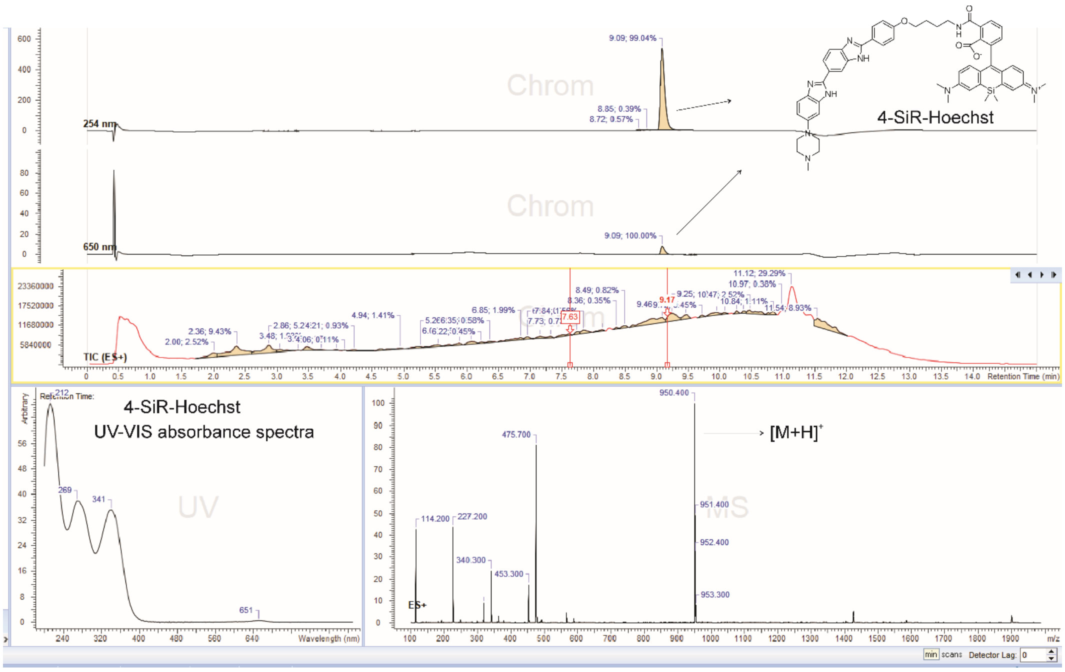
4-SiR-Hoechst
4-SiR-Hoechst
UV-VIS absorbance spectra
[M+H]+

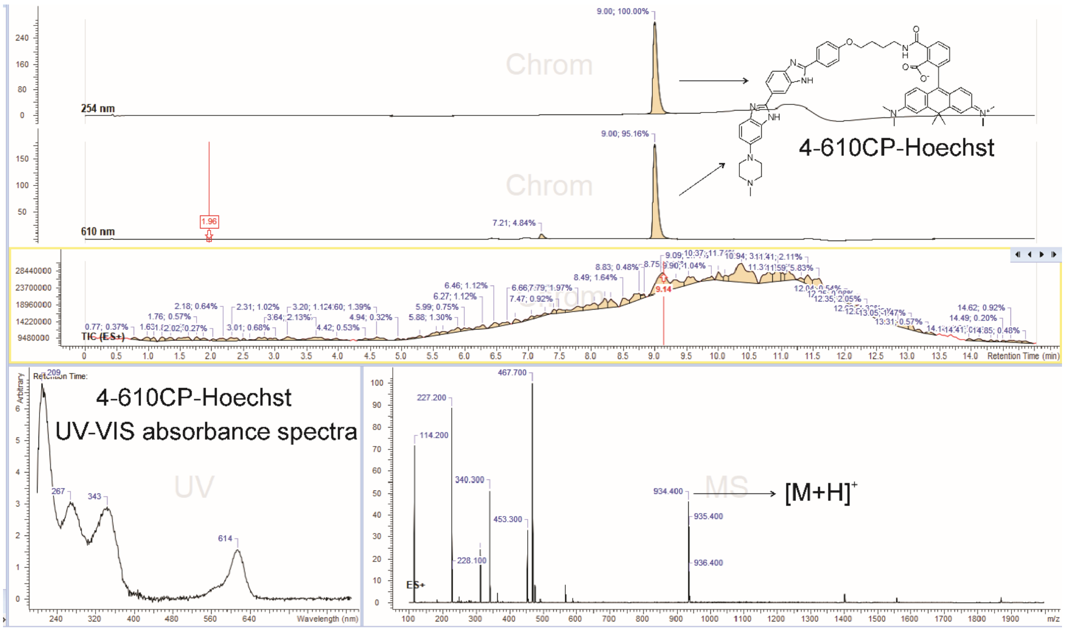
4-610CP-Hoechst
4-610CP-Hoechst
UV-VIS absorbance spectra
[M+H]+

### NMR spectra

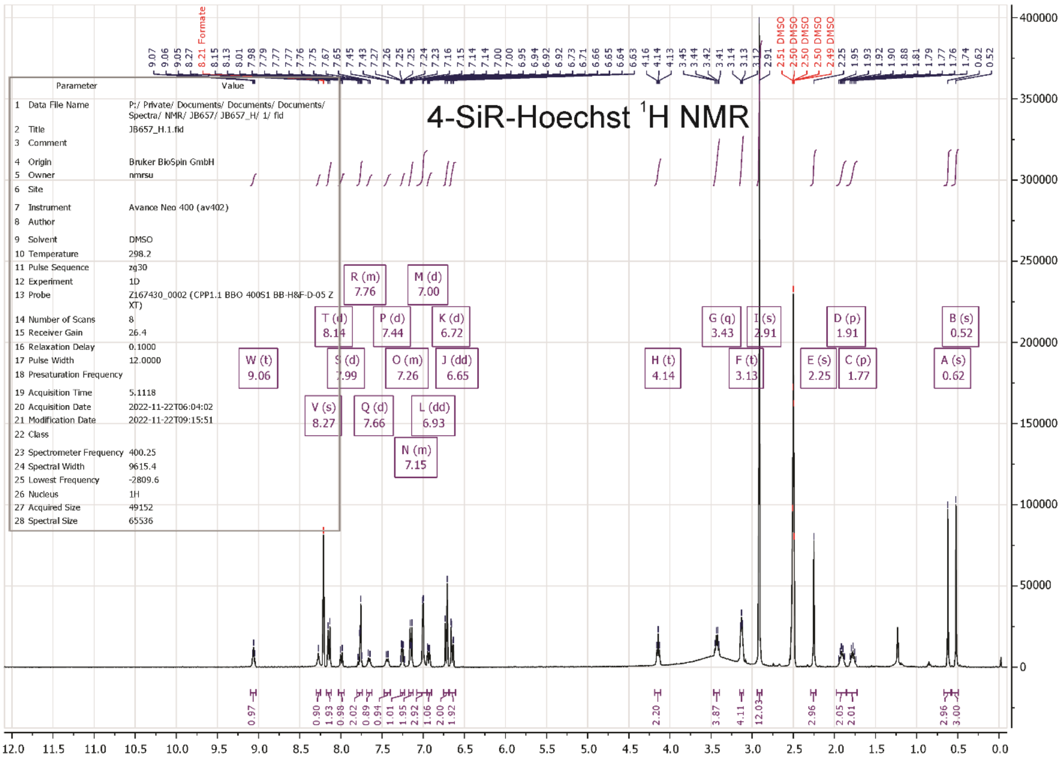
4-SiR-Hoechst 1H NMR
Parameter Value
1 Data File Name P:/ Private/ Documents/ Documents/ Documents/ Spectra/ NMR/ JB657/ JB657_H/ 1/ fid
2 Title JB657_H.1.fid
3 Comment
4 Origin Bruker BioSpin GmbH
5 Owner nmrsu
6 Site
7 Instrument Avance Neo 400 (av402)
8 Author
9 Solvent DMSO
10 Temperature 298.2
11 Pulse Sequence zg30
12 Experiment 1D
13 Probe Z167430_0002 (CPP1.1 BBO 400S1 BB-H&F-D-05 Z XT)
14 Number of Scans 8
15 Receiver Gain 26.4
16 Relaxation Delay 0.1000
17 Pulse Width 12.0000
18 Presaturation Frequency
19 Acquisition Time 5.1118
20 Acquisition Date 2022-11-22T06:04:02
21 Modification Date 2022-11-22T09:15:51
22 Class
23 Spectrometer Frequency 400.25
24 Spectral Width 9615.4
25 Lowest Frequency -2809.6
26 Nucleus 1H
27 Acquired Size 49152
28 Spectral Size 65536

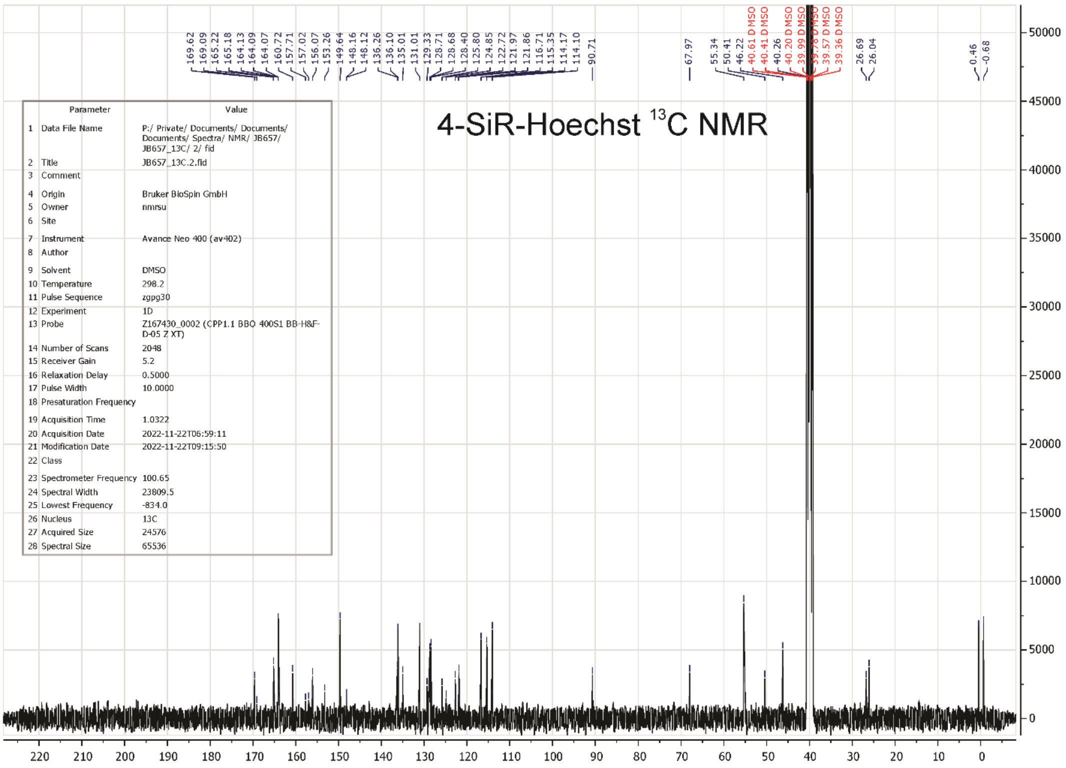
4-SiR-Hoechst 13C NMR
Parameter Value
1 Data File Name P:/ Private/ Documents/ Documents/ Documents/ Spectra/ NMR/ JB657/ JB657_13C/ 2/ fid
2 Title JB657_13C.2.fid
3 Comment
4 Origin Bruker BioSpin GmbH
5 Owner nmrsu
6 Site
7 Instrument Avance Neo 400 (av402)
8 Author
9 Solvent DMSO
10 Temperature 298.2
11 Pulse Sequence zgpg30
12 Experiment 1D
13 Probe Z167430_0002 (CPP1.1 BBO 400S1 BB-H&F-D-05 Z XT)
14 Number of Scans 2048
15 Receiver Gain 5.2
16 Relaxation Delay 0.5000
17 Pulse Width 10.0000
18 Presaturation Frequency
19 Acquisition Time 1.0322
20 Acquisition Date 2022-11-22T06:59:11
21 Modification Date 2022-11-22T09:15:50
22 Class
23 Spectrometer Frequency 100.65
24 Spectral Width 23809.5
25 Lowest Frequency -834.0
26 Nucleus 13C
27 Acquired Size 24576
28 Spectral Size 65536

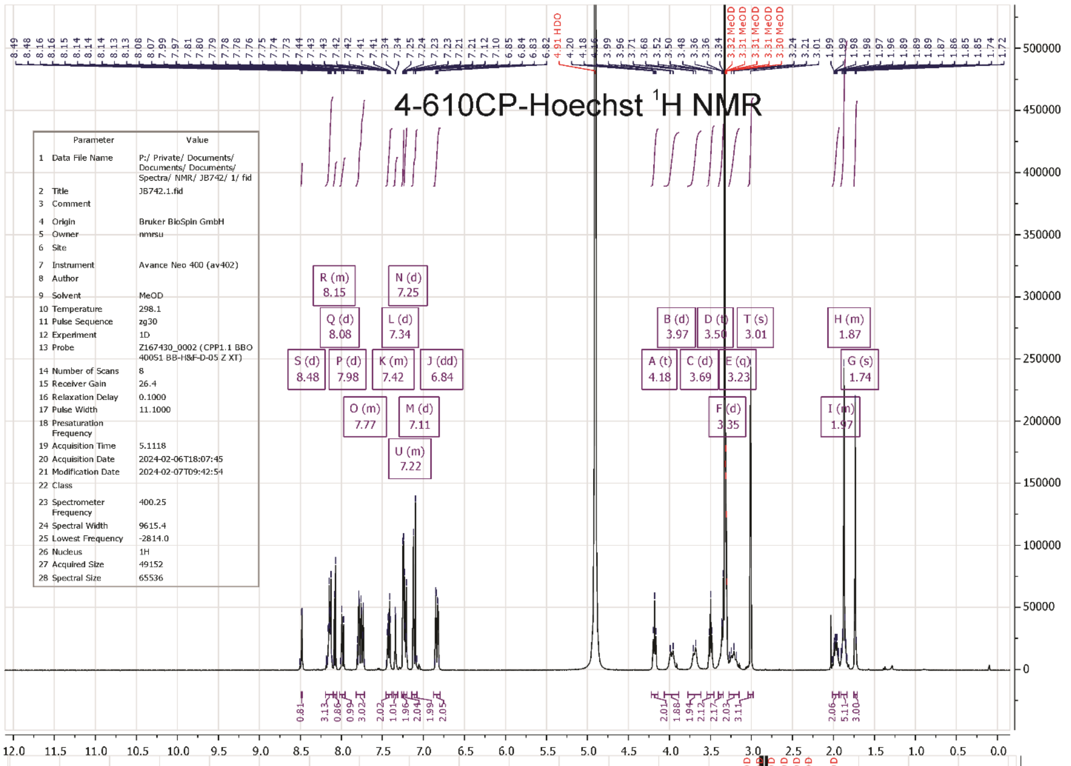
4-610CP-Hoechst $^{1}$H NMR

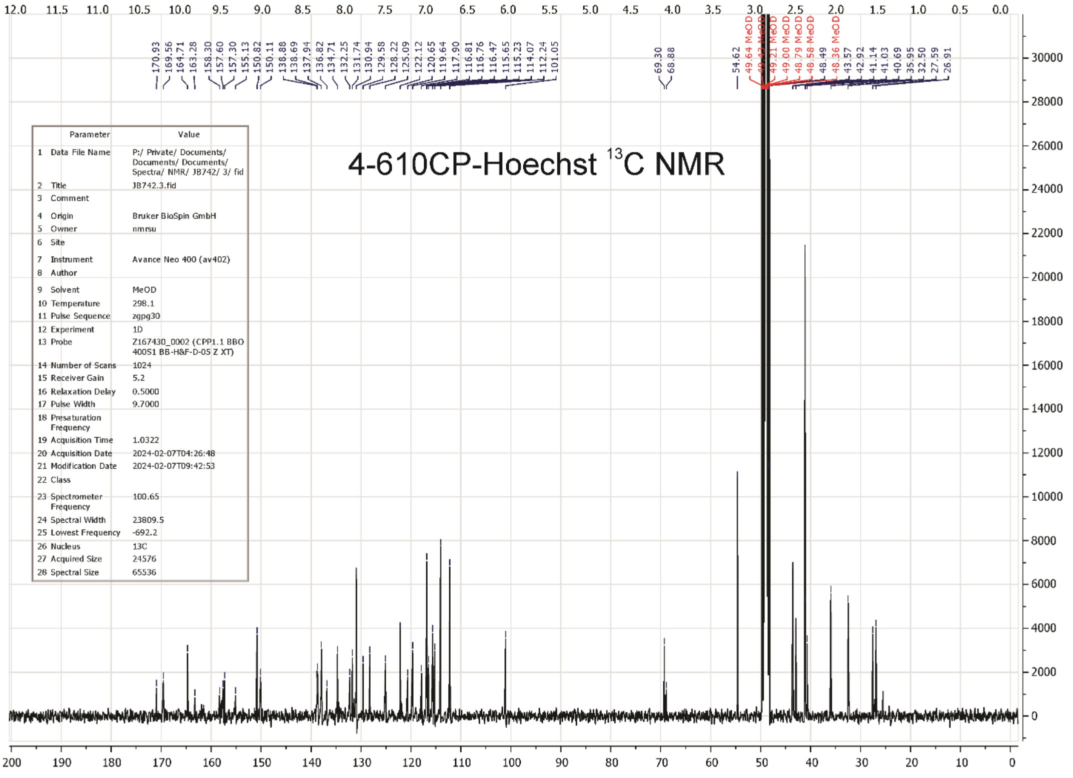
4-610CP-Hoechst $^{13}$C NMR

### Supplementary Tables

#### Table S1. Properties of free probes

| Probe name | In PBS | | | In PBS + 0.1% SDS | | | | Ref |
|---|---|---|---|---|---|---|---|---|
| | $\lambda_{max}^{Abs}$ (nm) | $\lambda_{max}^{em}$ (nm) | QY | $\lambda_{max}^{Abs}$ (nm) | $\lambda_{max}^{em}$ (nm) | QY | τ (ns), A (%) | |
| Hoechst 33342 | 340 | 510 | 0.026 ± 0.001 | 349 | 489 | 0.541± 0.002 | 2.59 (42%)<br>4.46 (58%) | [3] |
| **4-610CP-Hoechst** | 630 | 640 | 0.05 ± 0.02 | 618 | 640 | 0.67 ± 0.03 | 4.14 ± 0.03 | This work |
| 5-610CP-Hoechst | 619 | 643 | 0.052 ± 0.004 | 614 | 641 | 0.576 ± 0.004 | 4.22 | [3] |
| 6-610CP-Hoechst | 626 | 654 | 0.033 ± 0.003 | 615 | 642 | 0.614 ± 0.002 | 4.32 | [3] |
| **4-SiR-Hoechst** | 666 | 670 | 0.01 ± 0.03 | 655 | 671 | 0.59 ± 0.01 | 3.90 ± 0.02 | This work |
| 5-SiR-Hoechst | 659 | 671 | 0.007 ± 0.001 | 651 | 673 | 0.538 ± 0.004 | 3.93 | [3] |
| 6-SiR-Hoechst | 667 | 673 | 0.003 ± 0.002 | 652 | 676 | 0.524 ± 0.001 | 3.94 | [3] |

#### Table S2. Properties of probe-hpDNA complexes

| **Probe name** | $\lambda_{max}^{Abs}$ **(nm)** | $\lambda_{max}^{em}$ **(nm)** | **QY** | **τ (ns), A (%)** | **hpDNA binding affinity** | | **Ref** |
|---|---|---|---|---|---|---|---|
| | | | | | $K_{d1}$ **(μM)** | $K_{d2}$ **(μM)** | |
| Hoechst 33342 | 352 | 455 | 0.823 ± 0.004 | 2.79 ± 0.01 | 0.0026 ± 0.0004 | - | [3] |
| **4-610CP-Hoechst** | **615** | **638** | **0.39 ± 0.03** | **3.04 ± 0.02** | **1.3 ± 0.1** | - | This work |
| 5-610CP-Hoechst | 614 | 641 | 0.432 ± 0.001 | 3.45 ± 0.01 | 0.35 ± 0.02 | - | [3] |
| 6-610CP-Hoechst | 618 | 644 | 0.282 ± 0.005 | 1.78 (50%)<br>3.05 (50%) | 0.065 ± 0.007 | 4.4 ± 1.8 | [3] |
| **4-SiR-Hoechst** | **660** | **670** | **0.29 ± 0.11** | **2.78 ± 0.04** | **5.2 ± 0.9** | - | This work |
| 5-SiR-Hoechst | 651 | 672 | 0.374 ± 0.006 | 2.99 ± 0.01 | 4.8 ± 0.2 | - | [3] |
| 6-SiR-Hoechst | 654 | 677 | 0.156 ± 0.002 | 1.07 (53%)<br>2.72 (47%) | 8.5 ± 1.7 | - | [3] |

**Table S3. Properties of probe–mouse serum albumin complexes.**

| Probe name | $\lambda_{max}^{Abs}$ (nm) | $\lambda_{max}^{em}$ (nm) | QY | τ (ns), A (%) | MSA binding affinity | |
|---|---|---|---|---|---|---|
| | | | | | $K_{d1}$ (μM) | $K_{d2}$ (μM) |
| Hoechst 33342 | 345 | 460 | 0.033± 0.007 | 0.99 ± 0.01, 57<br>3.90 ± 0.02, 43 | 4 ± 7 | > 100 |
| 4-610CP-Hoechst | 624 | 640 | 0.02 ± 0.003 | 0.93 ± 0.13, 55<br>3.56 ± 0.10, 45 | 19 ± 15 | - |
| 5-610CP-Hoechst | 618 | 636 | 0.04 ± 0.02 | 0.53 ± 0.02, 55<br>3.69 ± 0.15, 45 | 14 ± 4 | - |
| 6-610CP-Hoechst | 616 | 634 | 0.07 ± 0.02 | 0.73 ± 0.06, 32<br>3.83 ± 0.07, 68 | 32 ± 16 | - |
| 4-SiR-Hoechst | 663 | 672 | 0.001 ± 0.004 | 0.79 ± 0.38, 34<br>2.93 ± 0.20, 66 | 96 ± 53 | - |
| 5-SiR-Hoechst | 661 | 666 | 0.014 ± 0.005 | 0.96 ± 0.05, 39<br>3.26 ± 0.03, 61 | 62 ± 14 | > 1000 |
| 6-SiR-Hoechst | 661 | 670 | 0.006 ± 0.003 | 1.34 ± 0.20, 28<br>3.56 ± 0.04, 72 | 62 ± 5 | - |

**Table S4. Properties of probe–human serum albumin complexes.**

| Probe name | $\lambda_{max}^{Abs}$ (nm) | $\lambda_{max}^{em}$ (nm) | QY | τ (ns), A (%) | HSA binding affinity | |
|---|---|---|---|---|---|---|
| | | | | | $K_{d1}$ (μM) | $K_{d2}$ (μM) |
| Hoechst 33342 | 350 | 460 | 0.082 ± 0.003 | 0.74 ± 0.02, 44<br>3.47 ± 0.01, 56 | 10 ± 4 | > 1000 |
| 4-610CP-Hoechst | 628 | 634 | 0.092 ± 0.002 | 1.02 ± 0.06, 35<br>3.77 ± 0.05, 65 | 57 ± 7 | - |
| 5-610CP-Hoechst | 616 | 632 | 0.172 ± 0.003 | 0.67 ± 0.55, 20<br>3.97 ± 0.44, 80 | 18 ± 4 | > 1000 |
| 6-610CP-Hoechst | 621 | 636 | 0.236 ± 0.009 | 0.72 ±0.07, 20<br>4.19 ±0.02, 80 | 31 ± 9 | > 1000 |
| 4-SiR-Hoechst | 674 | 674 | 0.003 ± 0.004 | - | 15 ± 9 | - |
| 5-SiR-Hoechst | 659 | 666 | 0.079 ± 0.007 | 0.34 ± 0.03, 26<br>3.73 ± 0.01, 74 | 25 ± 6 | > 1000 |
| 6-SiR-Hoechst | 663 | 674 | 0.060 ± 0.001 | 0.76 ± 0.28, 30<br>3.66 ± 0.04, 70 | 73 ± 59 | > 1000 |

**Table S5. Properties of probe–bovine serum albumin complexes.**

| Probe name | $\lambda_{max}^{Abs}$ (nm) | $\lambda_{max}^{em}$ (nm) | QY | τ (ns), A (%) | BSA binding affinity | |
|---|---|---|---|---|---|---|
| | | | | | $K_{d1}$ (μM) | $K_{d2}$ (μM) |
| Hoechst 33342 | 345 | 460 | 0.115 ± 0.003 | 0.74 ±0.03, 47<br>3.41 ±0.02, 53 | 48 ± 8 | - |
| 4-610CP-Hoechst | 627 | 630 | 0.210 ± 0.006 | 1.16 ± 0.08, 35<br>4.11 ±0.03, 65 | 132 ± 18 | - |
| 5-610CP-Hoechst | 618 | 634 | 0.106 ± 0.003 | 0.68 ± 0.18, 43<br>3.80 ± 0.05, 57 | 334 ± 70 | - |
| 6-610CP-Hoechst | 622 | 632 | 0.181 ± 0.007 | 0.88 ± 0.03, 45<br>3.94 ± 0.03, 55 | 136 ± 17 | - |
| 4-SiR-Hoechst | 669 | 672 | 0.027 ± 0.003 | - | 557 ± 520 | - |
| 5-SiR-Hoechst | 658 | 666 | 0.071 ± 0.004 | 0.89 ± 0.05, 37<br>3.76 ±0.02, 63 | 299 ± 77 | - |
| 6-SiR-Hoechst | 662 | 668 | 0.047 ± 0.002 | 0.66 ± 0.01, 66<br>3.45 ± 0.01, 34 | 361 ± 87 | - |

#### Table S6. Properties of dye-albumin complexes

| Dye | Serum albumin binding affinity | | | | | | | | |
|---|---|---|---|---|---|---|---|---|---|
| | Bovine serum albumin | | | Mouse serum albumin | | | Human serum albumin | | |
| | $\lambda_{max}^{Abs}$ (nm) | $\lambda_{max}^{em}$ (nm) | $K_d$ (μM) | $\lambda_{max}^{Abs}$ (nm) | $\lambda_{max}^{em}$ (nm) | $K_d$ (μM) | $\lambda_{max}^{Abs}$ (nm) | $\lambda_{max}^{em}$ (nm) | $K_d$ (μM) |
| 4-610CP-COOH | 615 | 632 | 573 ± 59 | 610 | 630 | 10 ± 2 | 620 | 630 | 46 ± 11 |
| 5-610CP-COOH | 610 | 630 | 430 ± 39 | 610 | 630 | 184 ± 114 | 620 | 630 | 59 ±10 |
| 6-610CP-COOH | 609 | 630 | 51 ± 5 | 610 | 630 | 126 ± 23 | 620 | 630 | 410 ± 150 |
| 4-SiR-COOH | 650 | 666 | 29 ± 3 | 640 | 664 | 0.16 ± 0.12 | 656 | 666 | 12 ± 1 |
| 5-SiR-COOH | 645 | 666 | 23 ± 5 | 640 | 664 | 17 ± 4 | 656 | 666 | 34 ± 3 |
| 6-SiR-COOH | 645 | 666 | 9.9 ± 0.5 | 640 | 664 | 5 ± 1 | 656 | 666 | 84 ± 9 |

#### Table S7. Laser power detected at different laser intensities in 2Photon microscopy

| Laser % | Laser power detected mW | |
|---|---|---|
| | At 800 nm | At 1300 nm |
| 100 | 145 | 38 |
| 50 | 73 | 19 |
| 25 | 36 | 9.3 |
| 12.5 | 18 | 4.4 |
| 6.5 | 9 | 2.2 |
| 3 | 4.2 | 1 |
| 1 | 1.4 | 0.3 |

**Table S8. List of mice used in the study**

| Sl.no | Probe injected | Animal No. | Gender | Weight (g) | Age in days |
|---|---|---|---|---|---|
| 1 | **4-610CP-Hoechst** | mouse no. 1 | Male | 47.8 | 146 |
| 2 | | mouse no. 2 | Female | 45.2 | 148 |
| 3 | | mouse no. 3 | Male | 45.68 | 160 |
| 4 | | mouse no. 4 | Female | 36.15 | 94 |
| 5 | | mouse no. 5 | Male | 48.85 | 97 |
| 6 | **5-610CP-Hoechst** | mouse no. 1 | Female | 44.9 | 367 |
| 7 | | mouse no. 2 | Male | 40.29 | 109 |
| 8 | | mouse no. 3 | Female | 38.6 | 115 |
| 9 | | mouse no. 4 | Female | 35.83 | 117 |
| 10 | | mouse no. 5 | Male | 37.56 | 102 |
| 11 | **6-610CP-Hoechst** | mouse no. 1 | Female | 45.79 | 132 |
| 12 | | mouse no. 2 | Female | 40.01 | 134 |
| 13 | | mouse no. 3 | Male | 43.1 | 141 |
| 14 | | mouse no. 4 | Male | 44.3 | 90 |
| 15 | | mouse no. 5 | Male | 38.3 | 90 |
| 16 | **4-SiR-Hoechst** | mouse no. 1 | Male | 45.8 | 167 |
| 17 | | mouse no. 2 | Male | 42.98 | 194 |
| 18 | | mouse no. 3 | Female | 32.08 | 174 |
| 19 | | mouse no. 4 | Male | 47.9 | 125 |
| 20 | | mouse no. 5 | Female | 44.38 | 203 |
| 21 | **5-SiR-Hoechst** | mouse no. 1 | Male | 37.9 | 102 |
| 22 | | mouse no. 2 | Female | 40.2 | 127 |
| 23 | | mouse no. 3 | Female | 43.4 | 129 |
| 24 | | mouse no. 4 | Male | 43.3 | 135 |
| 25 | | mouse no. 5 | Female | 43.05 | 142 |
| 26 | **6-SiR-Hoechst** | mouse no. 1 | Male | 45.19 | 169 |
| 27 | | mouse no. 2 | Female | 43.04 | 169 |
| 28 | | mouse no. 3 | Male | 49.06 | 174 |
| 29 | | mouse no. 4 | Female | 37.5 | 513 |
| 30 | | mouse no. 5 | Male | 37.89 | 91 |

**Table S9. Constituents of the intravenous injection solution injected to mice**

| Investigated Probe | Probe in DMSO (µL) | NaCl (µL) | Pluronic acid (µL) | Total injection volume (µL) |
|---|---|---|---|---|
| 4/5/6-610CP Hoechst | 20 | 100 | 20 | 140 |
| 4/5/6-SiR Hoechst | 20 | 100 | 20 | 140 |

**Table S10. Imaging parameters for confocal microscopy of mice tissues.**

| Probe injected | Organ | Laser (nm) | Laser power (%) | Gain (%) | Detection (nm) |
|---|---|---|---|---|---|
| **4-610CP-Hoechst** | Liver | 633 | 2.2 - 4.2 | 190 - 499.8 | 642 - 794 |
| | Kidney | 633 | 1.25 - 4.21 | 115.1 - 309.1 | 642 - 794 |
| | Heart | 633 | 4.21 - 14.8 | 222.2 - 500 | 642 - 794 |
| | Lungs | 633 | 2.16 - 4.35 | 222.5 - 409.3 | 642 - 794 |
| | Brain | 633 | 4.3 - 41.9 | 222.5 - 500 | 642 - 794 |
| **5-610CP-Hoechst** | Liver | 633 | 0.7 - 5.2 | 10 - 85.9 | 600 - 700 |
| | Kidney | 633 | 0.3 - 6.0 | 10 - 165.4 | 600 - 700 |
| | Heart | 633 | 0.7 - 6.0 | 54.2 - 344.8 | 600 - 700 |
| | Lungs | 633 | 0.7 - 6.0 | 10 - 344.8 | 600 - 700 |
| | Brain | 633 | 0.4 - 17.5 | 115.1 - 500 | 600 - 700 |
| **6-610CP-Hoechst** | Liver | 633 | 1.8 - 17.5 | 11.5 - 452.6 | 610 - 794 |
| | Kidney | 633 | 1.8 - 17.5 | 20.4 - 452.6 | 610 - 794 |
| | Heart | 633 | 1.8 - 20.8 | 80.3 - 497.9 | 610 - 794 |
| | Lungs | 633 | 3.6 - 28.1 | 12.5 - 500 | 610 - 794 |
| | Brain | 633 | 2.5 - 28.1 | 31.7 - 462.4 | 610 - 794 |
| **4-SiR-Hoechst** | Liver | 633 | 10.26 - 25.76 | 68.2 - 386.1 | 642 - 794 |
| | Kidney | 633 | 23.87 - 25.7 | 227.8 - 386.1 | 642 - 794 |
| | Heart | 633 | 17.5 - 25.8 | 376.3 - 407 | 642 - 794 |
| | Lungs | 633 | 17.5 - 25.7 | 289.1 - 407 | 642 - 794 |
| | Brain | 633 | 10.2 - 22.7 | 53.2 - 386.1 | 642 - 794 |
| **5-SiR-Hoechst** | Liver | 633 | 1.46 - 7.0 | 40.2 - 315 | 610 - 795 |
| | Kidney | 633 | 1.46 - 7.0 | 36.7 - 467.2 | 610 - 795 |
| | Heart | 633 | 3.4 - 17.5 | 83.8 - 500 | 610 - 795 |
| | Lungs | 633 | 3.4 - 7.0 | 117.4 - 444.5 | 610 - 795 |
| | Brain | 633 | 4.8 - 25.6 | 63.8 - 500 | 610 - 795 |
| **6-SiR-Hoechst** | Liver | 633 | 3.73 - 15.5 | 142.3 - 246.6 | 642 - 794 |
| | Kidney | 633 | 3.73 - 20.07 | 230.1 - 433 | 642 - 794 |
| | Heart | 633 | 10.9 - 25.5 | 362.3 - 447.8 | 642 - 794 |
| | Lungs | 633 | 10.9 - 21.8 | 279.9 - 407.6 | 642 - 794 |
| | Brain | 633 | 9.1 - 22.7 | 132 - 499.6 | 642 - 794 |

**Table S11. Imaging parameters for 2-photon microscopy of mice tissues.**

| Probe injected | Organ | Laser (%) at 800 nm* and 1300 nm** | Gain (%) | Detection (nm) |
|---|---|---|---|---|
| **4-610CP-Hoechst** | Liver | 2.6 | 58-80 | 590-650 |
| | Kidney | 2-4.9 | 60-80 | |
| | Heart | 4.3-17.25 | 60-80 | |
| | Lungs | 2.25-7.5 | 60-80 | |
| | Brain | 5-11.5 | 60-82 | |
| **5-610CP-Hoechst** | Liver | 1.55-3.8 | 58-72 | |
| | Kidney | 2.05-2.6 | 65-70 | |
| | Heart | 2.7-4.35 | 60-76 | |
| | Lungs | 1.95-3.85 | 60-70 | |
| | Brain | 4 | 4 | |
| **6-610CP-Hoechst** | Liver | 1.9-6 | 60-72 | |
| | Kidney | 0.9-12.25 | 61-72 | |
| | Heart | 8.5-10.25 | 70-72 | |
| | Lungs | 4-9 | 60-72 | |
| | Brain | 5 | 5 | |
| **4-SiR-Hoechst** | Liver | 53-70 | 80 | 647LP (≥647) |
| | Kidney | 58-80 | 80 | |
| | Heart | 57-75 | 80-82 | |
| | Lungs | 44-65 | 80-81 | |
| | Brain | 58-78 | 80-81 | |
| **5-SiR-Hoechst** | Liver | 13.5-23.25 | 78-80 | |
| | Kidney | 16.5-17.5 | 80 | |
| | Heart | 39-57 | 70-80 | |
| | Lungs | 13.5-20.75 | 70-80 | |
| | Brain | 77 | 78 | |
| **6-SiR-Hoechst** | Liver | 59-70 | 80 | |
| | Kidney | 61-70 | 80 | |
| | Heart | 61-81 | 80-82 | |
| | Lungs | 81 | 80 | |
| | Brain | 71-78 | 80 | |

### Supplementary Figures

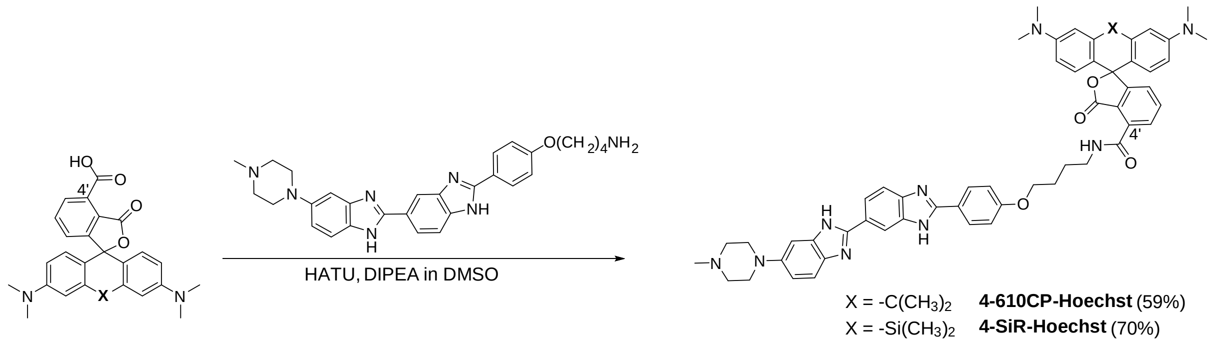

**Figure S1. Synthesis of 4' regioisomer rhodamine–Hoechst probes.**

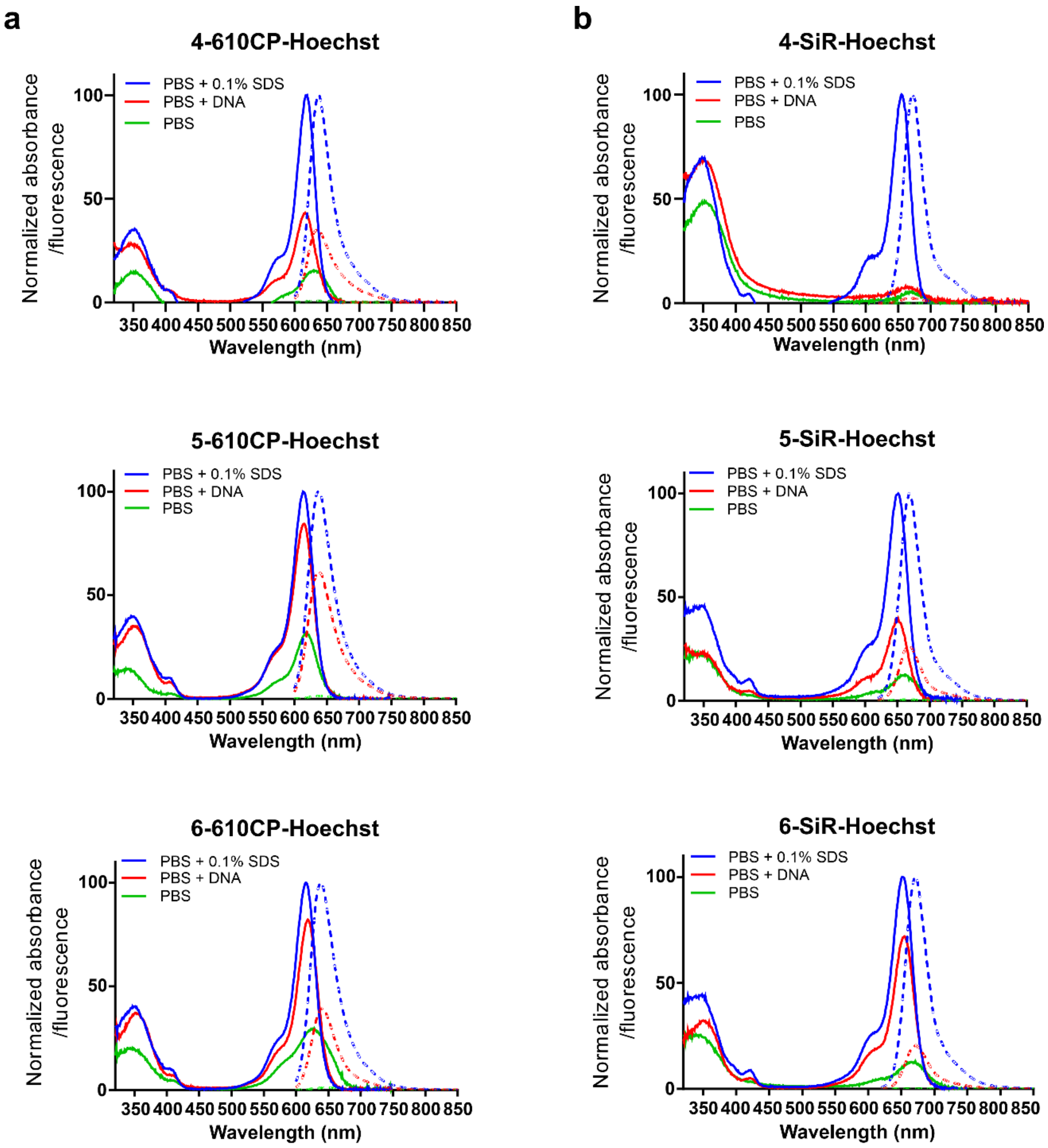

**Figure S2. Spectral properties of DNA probes.**

Absorbance (solid line) and fluorescence (dotted line) spectra of Rhodamine-Hoechst probes: 4-, 5-, 6-610CP-Hoechst (**a** column) and 4-, 5-, 6-SiR-Hoechst (**b** column) in PBS (green), PBS containing 30 µM hpDNA oligonucleotide (red) and PBS containing 0.1% SDS (blue). 2 µM probes were incubated at RT for 2 h before measurements. Spectra are normalized to the samples containing 0.1% SDS and presented as averages of three independent experiments.

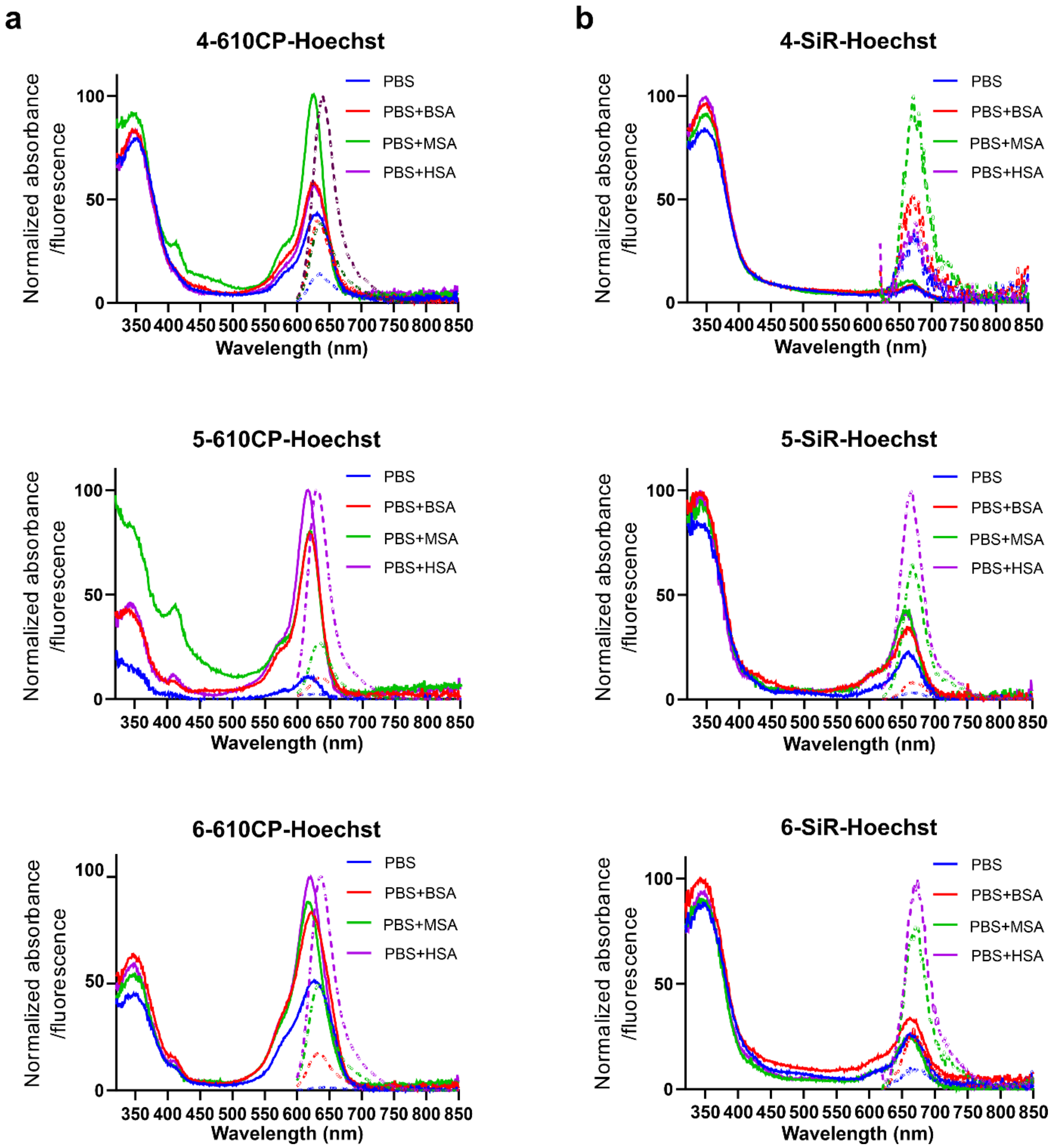

**Figure S3. Spectral properties of DNA probes when interacting with serum albumin.**

Absorbance (solid line) and fluorescence (dotted line) spectra of Rhodamine-Hoechst probes: 4-, 5-, 6-610CP-Hoechst (**a** column) and 4-, 5-, 6-SiR-Hoechst (**b** column) in PBS (blue), PBS containing BSA (red), MSA (green) and HSA (magenta). 1μM probes were incubated at RT for 2 h before measurements. Spectra are normalized to the samples containing the highest signal and presented as averages of three independent experiments.

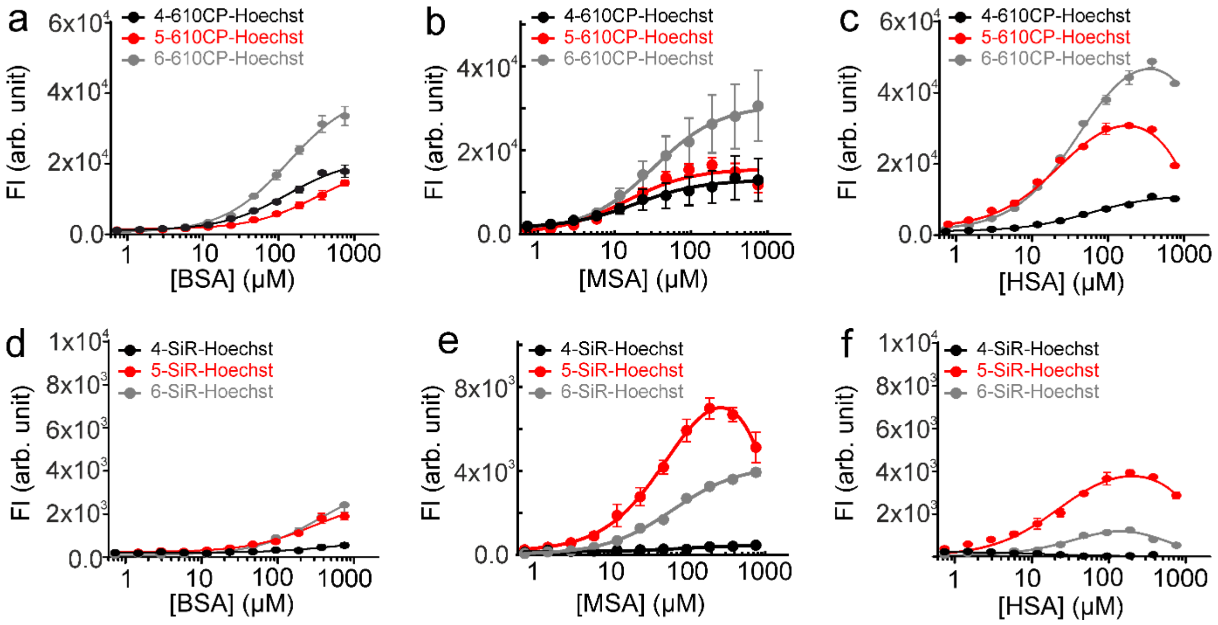

**Figure S4. Interaction of DNA probes with serum albumin.**

Titration of 200 nM probes: 4-, 5-, 6-610CP-Hoechst (**a, b, c**) and 4-, 5-, 6-SiR-Hoechst (**d, e, f**) with purified BSA (**a, d**), MSA (**b, e**) and HSA (**c, f**). Two site-binding equation is fitted to the data points for 5-, 6- 610CP-Hoechst and SiR-Hoechst with purified HSA and also for 5-SiR-Hoechst with purified MSA. A single site binding equation was fitted to the others. Data are presented as mean ± s. e. m. of n=3 experimental replicates. BSA - bovine serum albumin. HSA - human serum; MSA - mouse serum albumin.

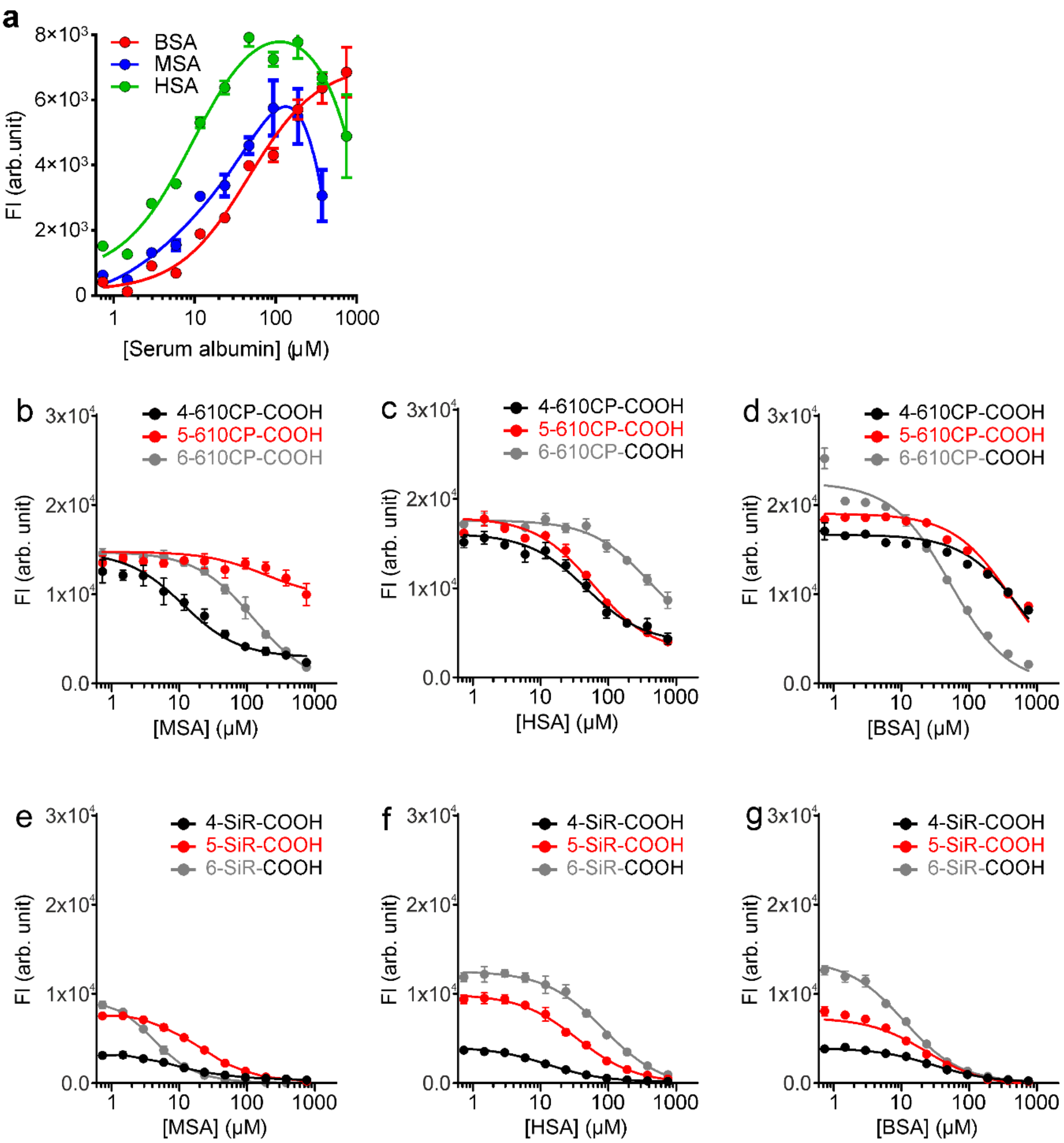

**Figure S5. Interaction of Hoechst 33342 and rhodamine-dyes with serum albumin.**

Titration of 0.2 µM Hoechst 33342 (**a**), 2 µM 4-, 5-, 6-610CP-COOH (**b, c, d**) or 2 µM 4-, 5-, 6-SiR-COOH (**e, f, g**) dyes with purified serum albumins - MSA, HSA or BSA. Hoechst 33342 – HSA and MSA data points were fitted to two site-binding equation. All other data points were fitted using a single site binding equation. Data are presented as mean ± s.e.m of n=3 experimental replicates. BSA, bovine serum albumin; FI, fluorescence signal; HSA, human serum; MSA, mouse serum albumin.

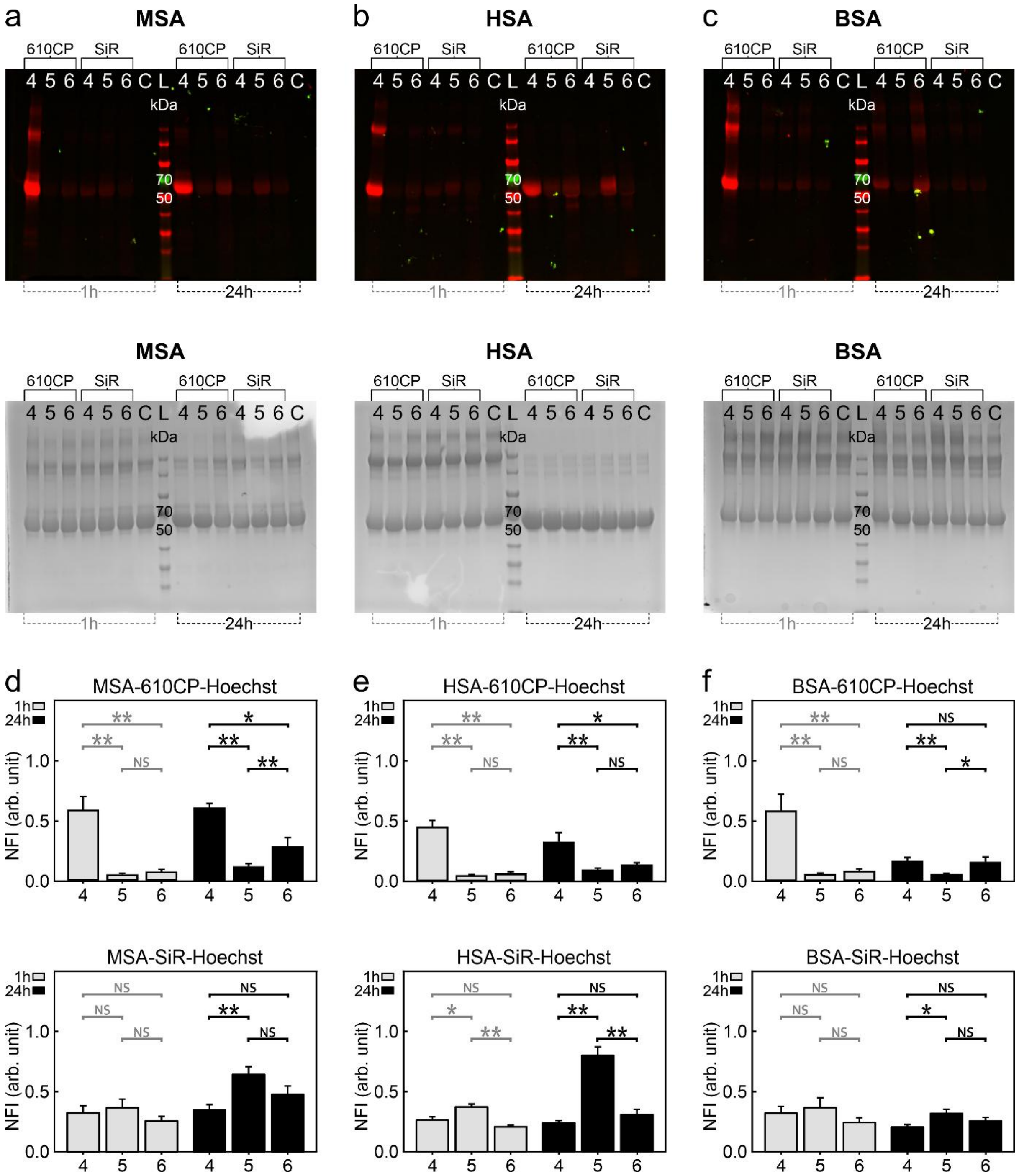

**Figure S6. Covalent interaction of purified mouse/human/bovine serum albumin with DNA probes.**

Representative SDS-PAGE gels showing probes: 4-, 5-, 6-610CP-Hoechst and 4-, 5-, 6-SiR-Hoechst covalently bonded to purified MSA (**a**), HSA (**b**), and BSA (**c**). 1 µM probes were incubated with purified MSA/HSA/BSA for 1 h and 24 h, and subsequently subjected to denaturation at 95˚C for 5 min followed by SDS-PAGE analysis. The gels were subjected to Coomassie blue staining (**a-c** lower row). In-gel fluorescence was quantified for crosslink formation between serum albumins and 610CP-Hoechst (**d-e** upper row) or SiR-Hoechst (**d**–**e** lower row) probes. The bar graphs showing the normalized fluorescence intensity of 66kDa bands on the gels. Data are mean ± s.d of n=5 experimental replicates; * p≤0.03, ** p≤0.002, NS-not significant in Mann Whitney t-test. Arb., arbitrary; BSA, bovine serum albumin; FS, fluorescence signal; HSA, human serum albumin; MSA, mouse serum albumin; NFI, normalized fluorescence intensity.

**a**

```
                          10        20        30        40        50        60        70        80
                  ....|....|....|....|....|....|....|....|....|....|....|....|....|....|....|....|
BSA_(CAA76847)  1   MKWVTFISLLLLFSSAYSRGVFRRDTHKSEIAHRFKDLGEEHFKGLVLIAFSQYLQQCPFDEHVKLVNELTEFAKTCVAD
HSA_(AAH36003)  1   MKWVTFISLLFLFSSAYSRGVFRRDAHKSEVAHRFKDLGEENFKALVLIAFAQYLQQCPFEDHVKLVNEVTEFAKTCVAD
MSA_(CAD29888)  1   MKWVTFLLLLFVSGSAFSRGVFRREAHKSEIAHRYNDLGEQHFKGLVLIAFSQYLQKCSYDEHAKLVQEVTDFAKTCVAD

                          90       100       110       120       130       140       150       160
                  ....|....|....|....|....|....|....|....|....|....|....|....|....|....|....|....|
BSA_(CAA76847)  81  ESHAGCEKSLHTLFGDELCKVASLRETYGDMADCCEKQEPERNECFLSHKDDSPDLPKLK.PDPNTLCDEFKADEKKFWG
HSA_(AAH36003)  81  ESAENCDKSLHTLFGDKLCTVATLRETYGEMADCCAKQEPERNECFLQHKDDNPNLPRLVRPEVDVMCTAFHDNEETFLK
MSA_(CAD29888)  81  ESAANCDKSLHTLFGDKLCAIPNLRENYGELADCCTKQEPERNECFLQHKDDNPSLPPFERPEAEAMCTSFKENPTTFMG

                         170       180       190       200       210       220       230       240
                  ....|....|....|....|....|....|....|....|....|....|....|....|....|....|....|....|
BSA_(CAA76847)  160 KYLYEIARRHPYFYAPELLYYANKYNGVFQECCQAEDKGACLLPKIETMREKVLTSSARQRLRCASIQKFGERALKAWSV
HSA_(AAH36003)  161 KYLYEIARRHPYFYAPELLFFAKRYKAAFTECCQAADKAACLLPKLDELRDEGKASSAKQRLKCASLQKFGERAFKAWAV
MSA_(CAD29888)  161 HYLHEVARRHPYFYAPELLYYAEQYNEILTQCCAEADKESCLTPKLDGVKEKALVSSVRQRMKCSSMQKFGERAFKAWAV

                         250       260       270       280       290       300       310       320
                  ....|....|....|....|....|....|....|....|....|....|....|....|....|....|....|....|
BSA_(CAA76847)  240 ARLSQKFPKAEFVEVTKLVTDLTKVHKECCHGDLLECADDRADLAKYICDNQDTISSKLKECCDKPLLEKSHCIAEVEKD
HSA_(AAH36003)  241 ARLSQRFPKAEFAEVSKLVTDLTKVHTECCHGDLLECADDRADLAKYICENQDSISSKLKECCEKPLLEKSHCIAEVEND
MSA_(CAD29888)  241 ARLSQTFPNADFAEITKLATDLTKVNKECCHGDLLECADDRAELAKYMCENQATISSKLQTCCDKPLLKKAHCLSEVEHD

                         330       340       350       360       370       380       390       400
                  ....|....|....|....|....|....|....|....|....|....|....|....|....|....|....|....|
BSA_(CAA76847)  320 AIPENLPPLTADFAEDKDVCKNYQEAKDAFLGSFLYEYSRRHPEYAVSVLLRLAKEYEATLEECCAKDDPHACYSTVFDK
HSA_(AAH36003)  321 EMPADLPSLAADFVESKDVCKNYAEAKDVFLGMFLYEYARRHPDYSVVLLLRLAKTYETTLEKCCAAADPHECYAKVFDE
MSA_(CAD29888)  321 TMPADLPAIAADFVEDQEVCKNYAEAKDVFLGTFLYEYSRRHPDYSVSLLLRLAKKYEATLEKCCAEANPPACYGTVLAE

                         410       420       430       440       450       460       470       480
                  ....|....|....|....|....|....|....|....|....|....|....|....|....|....|....|....|
BSA_(CAA76847)  400 LKHLVDEPQNLIKQNCDQFEKLGEYGFQNALIVRYTRKVPQVSTPTLVEVSRSLGKVGTRCCTKPESERMPCTEDYLSLI
HSA_(AAH36003)  401 FKPLVEEPQNLIKQNCELFEQLGEYKFQNALLVRYTKKVPQVSTPTLVEVSRNLGKVGSKCCKHPEAKRMPCAEDYLSVV
MSA_(CAD29888)  401 FQPLVEEPKNLVKTNCDLYEKLGEYGFQNAILVRYTQKAPQVSTPTLVEAARNLGRVGTKCCTLPEDQRLPCVEDYLSAI

                         490       500       510       520       530       540       550       560
                  ....|....|....|....|....|....|....|....|....|....|....|....|....|....|....|....|
BSA_(CAA76847)  480 LNRLCVLHEKTPVSEKVTKCCTESLVNRRPCFSALTPDETYVPKAFDEKLFTFHADICTLPDTEKQIKKQTALVELLKHK
HSA_(AAH36003)  481 LNQLCVLHEKTPVSDRVTKCCTESLVNRRPCFSALEVDETYVPKEFNAETFTFHADICTLSEKERQIKKQTALVELVKHK
MSA_(CAD29888)  481 LNRVCLLHEKTPVSEHVTKCCSGSLVERRPCFSALTVDETYVPKEFKAETFTFHSDICTLPEKEKQIKKQTALAELVKHK

                         570       580       590       600
                  ....|....|....|....|....|....|....|....
BSA_(CAA76847)  560 PKATEEQLKTVMENFVAFVDKCCAADDKEACFAVEGPKLVVSTQTALA.
HSA_(AAH36003)  561 PKATKEQLKAVMDDFAAFVEKCCKADDKETCFAEEGKKLVAASQAALGL
MSA_(CAD29888)  561 PKATAEQLKTVMDDFAQFLDTCCKAADKDTCFSTEGPNLVTRCKDALA.
```

**b**

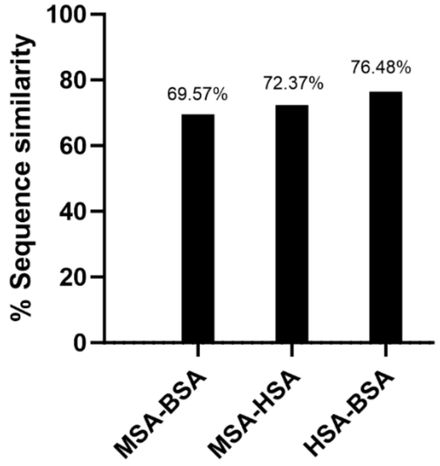

**Figure S7: Multiple alignment of MSA, HSA, and BSA sequence.**

**a** Amino acid sequences of MSA (P07724), HSA (P02768), and BSA (P02769) aligned using Multalin version 5.4.1. Regions interacting with rhodamine-Hoechst ligands are highlighted: magenta – SiR-based ligands, green – 610CP-based ligands and cyan – both types of ligands. **b** Bar graph representing percentages of sequence similarities between MSA, HSA and BSA.

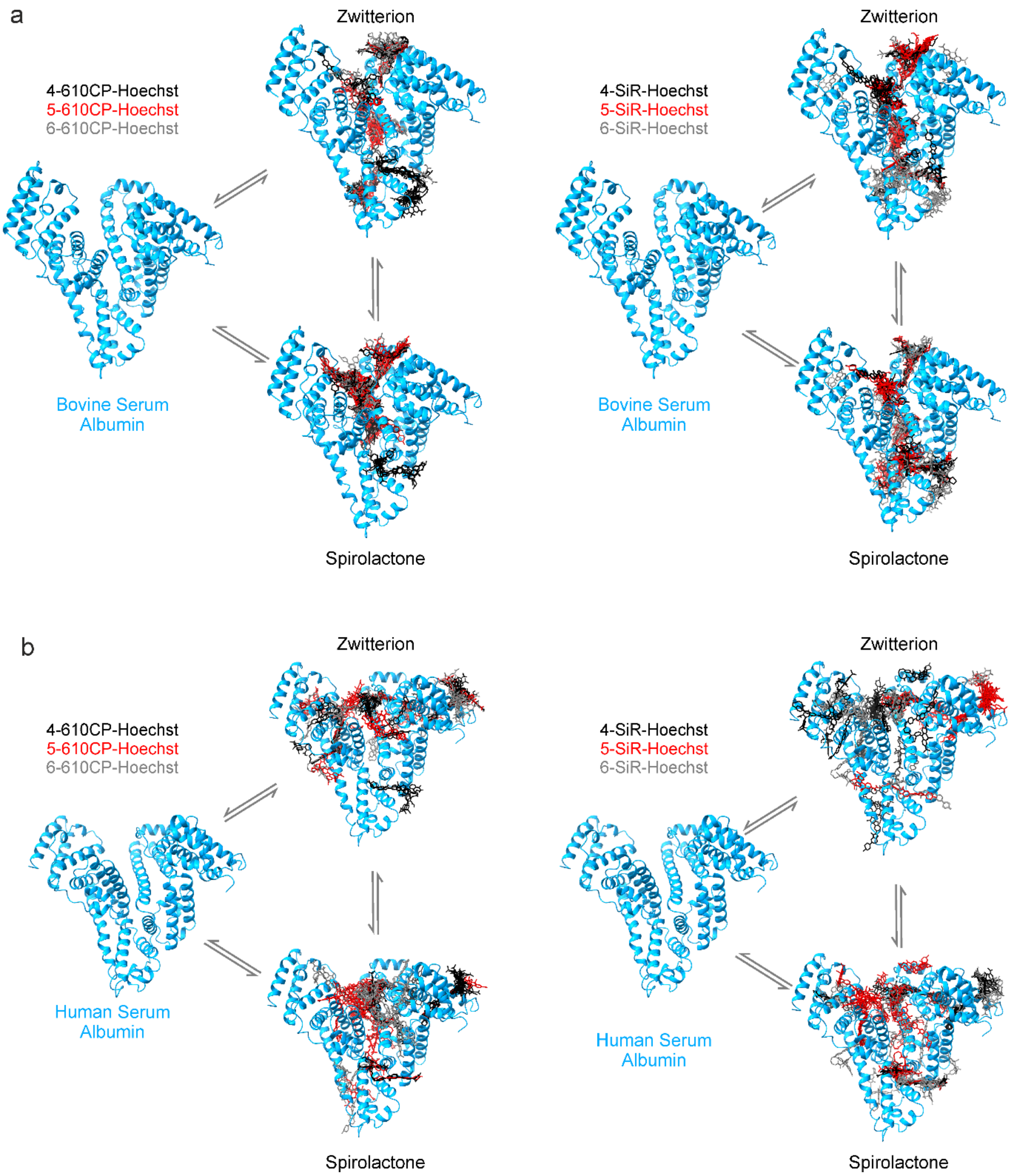

**Figure S8. Modelling of Interaction of rhodamine-Hoechst probes with bovine (a) or human (b) serum albumins.**

Molecular docking model of bovine or human serum albumin bound to all three regioisomers of 610CP-Hoechst and SiR-Hoechst. Two forms of the DNA probes are shown: “open” (zwitterion) and “closed” (spirolactone). Apo-protein structures were taken from the following X-ray structures: human - PDB ID 4BKE[4] and bovine PDB ID 4JK4 were used.

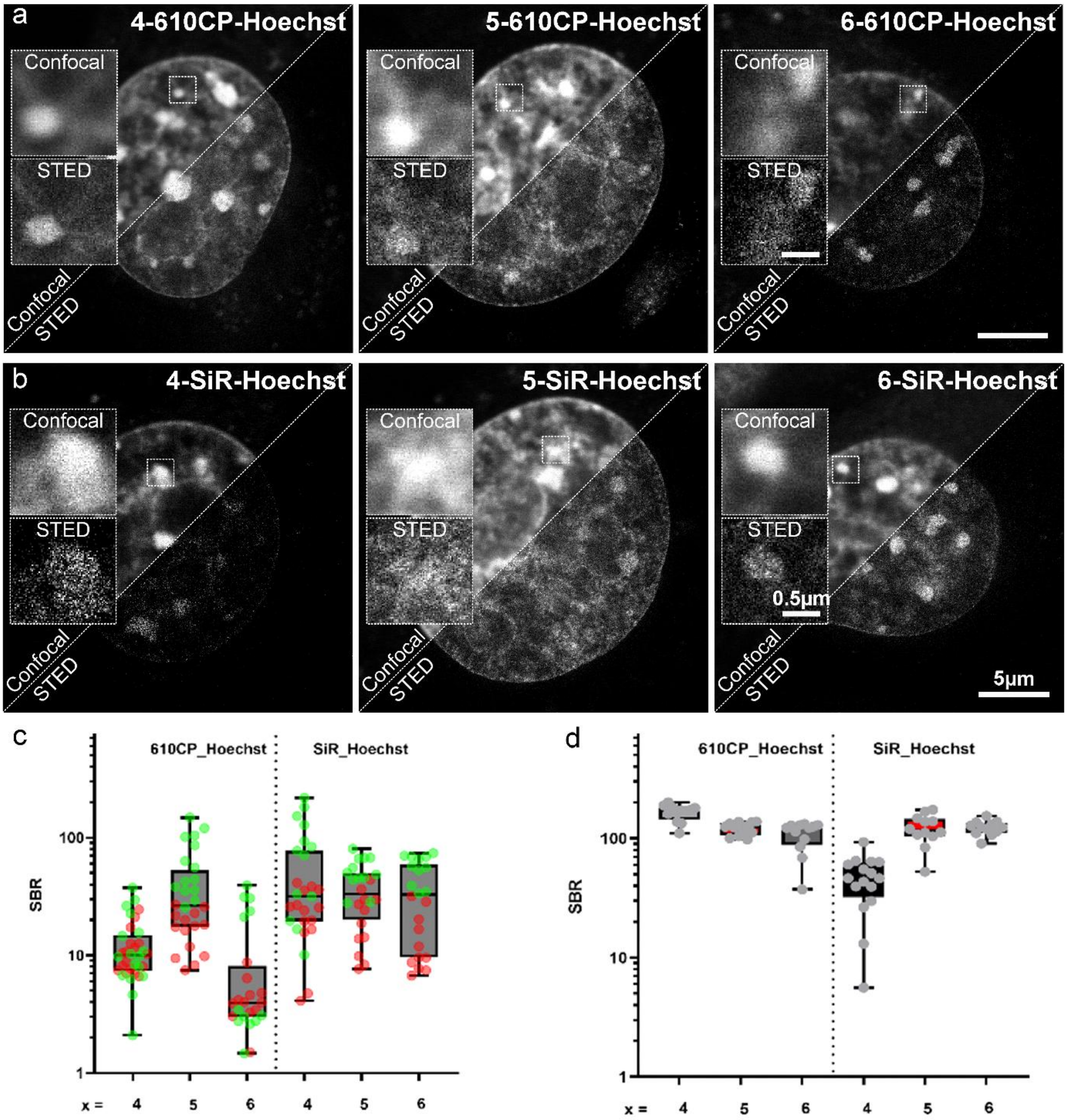

**Figure S9. Staining efficiency of rhodamine-Hoechst probes in living NIH3T3 mouse fibroblasts.**

Confocal and STED imaging of nuclei in living NIH3T3 mouse fibroblasts stained with 200 nM Rhodamine-Hoechst derivatives: 4-, 5-, 6-610CP-Hoechst (**a**) and 4-, 5-, 6-SiR-Hoechst (**b**). Images were acquired on Abberior Expert Line nanoscope. Scale bar: 5 µm. **c** Data points are signal to background ratios (SBRs) for 3 nuclei (red for confocal and green for STED). **d** Data points are fluorescence intensity signal from nucleoli in each nucleus (3 nuclei) from STED images. Numbers: 4, 5 and 6 indicate different regioisomer.

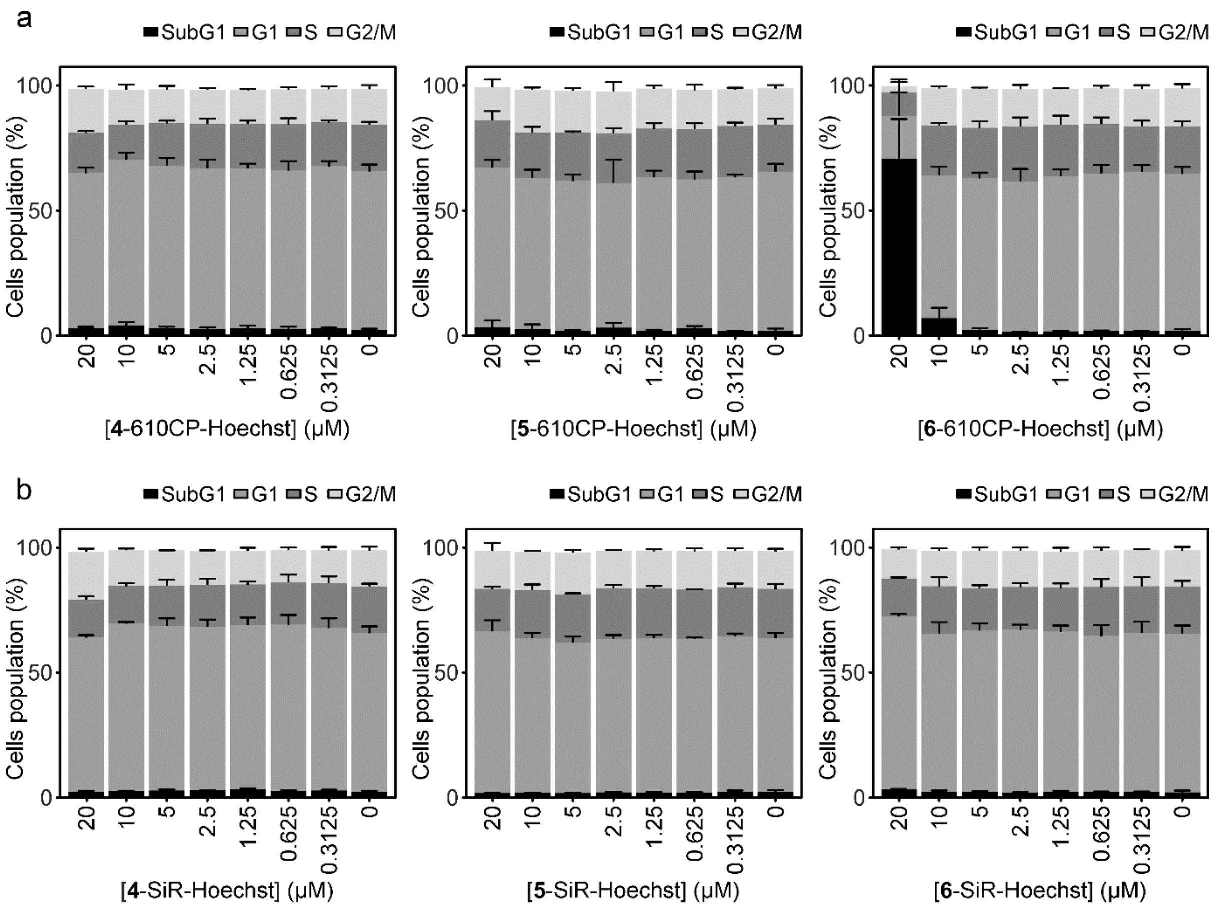

**Figure S10. Cell cytotoxicity induced by fluorescent DNA probes.**

NIH3T3 mouse fibroblasts were incubated with the indicated concentrations of the 4-, 5-, 6-610CP-Hoechst (**a**) and 4-, 5-, 6-SiR-Hoechst (**b**) DNA probes at 37 °C for 24 h in a humidified 5% $CO_2$ incubator. The bar graphs indicate cell populations at given cell cycle phases (Sub G1, G1, S and G2-M), identified by the amount of DNA in the investigated cells. Experimental data are averages of n=3 independent experiments and presented as mean ± SD.

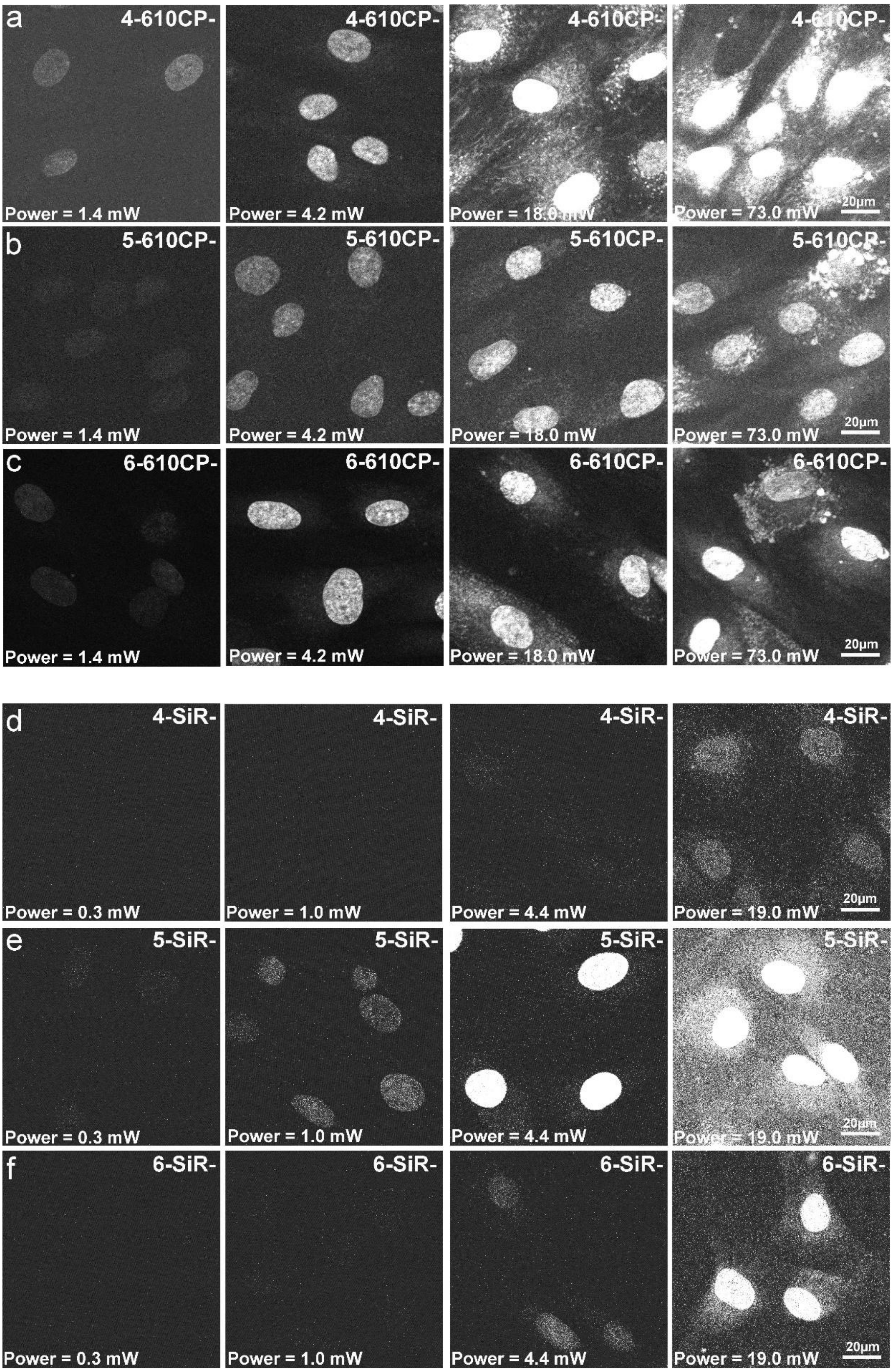

**Figure S11. The two-photon microscopy of human fibroblast cells stained by DNA probes.**

Human fibroblast cells stained with 200 nM of (**a** - **c**) 4-, 5-, 6-610CP-Hoechst and (**d** - **f**) 4-, 5-, 6-SiR-Hoechst for 1 hr prior to imaging. The images were captured at different laser powers as indicated on the image. Scale bar 20 μm.

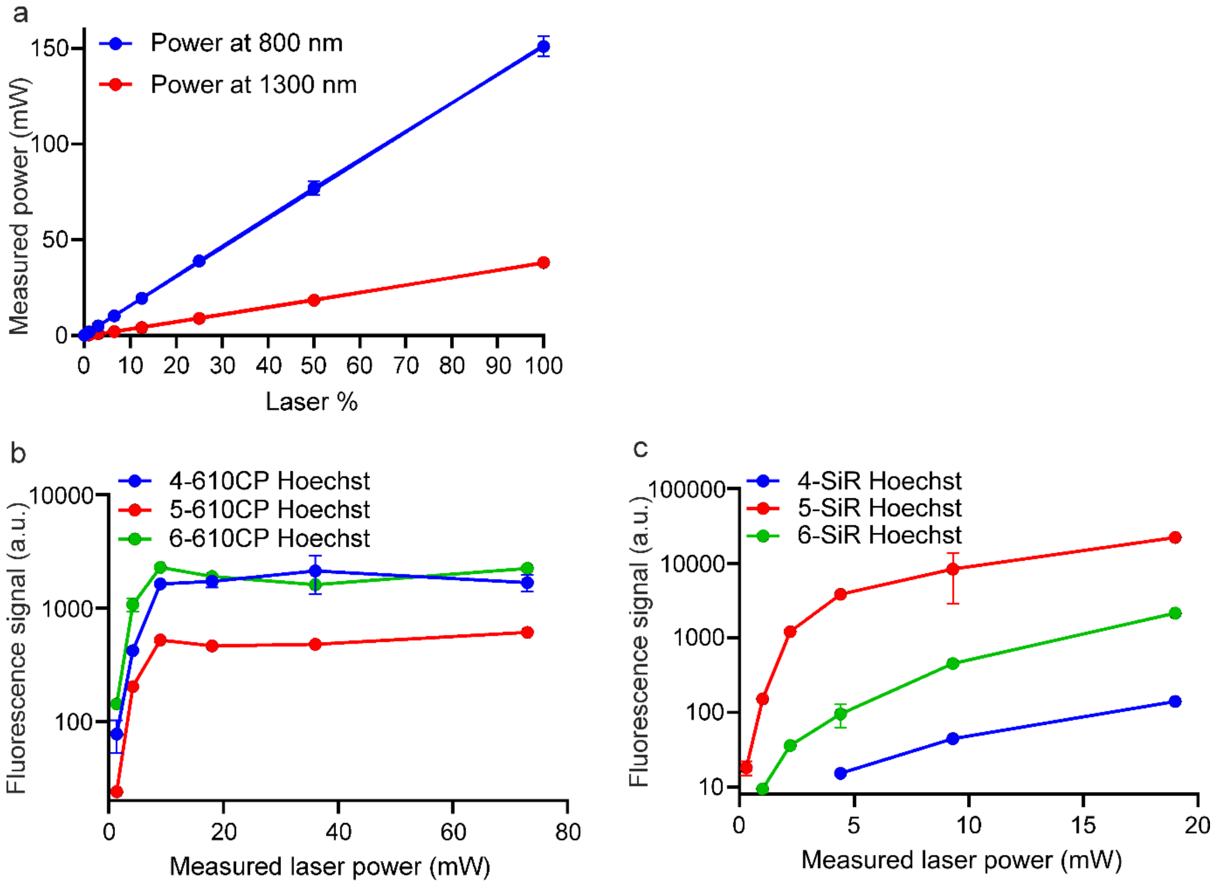

**Figure S12. Plots of laser power measurements in 2Photon microscopy.**

**a** Laser % vs measured power plots recorded at 800 nm and 1300 nm. The fluorescence intensity of the nuclei is plotted against the laser power for cells stained with 610CP-Hoechst (**b**) and SiR-Hoechst (**c**). Data points are fluorescence intensity signal of nuclei from 3 images (n=3) and are presented as mean ± SEM.

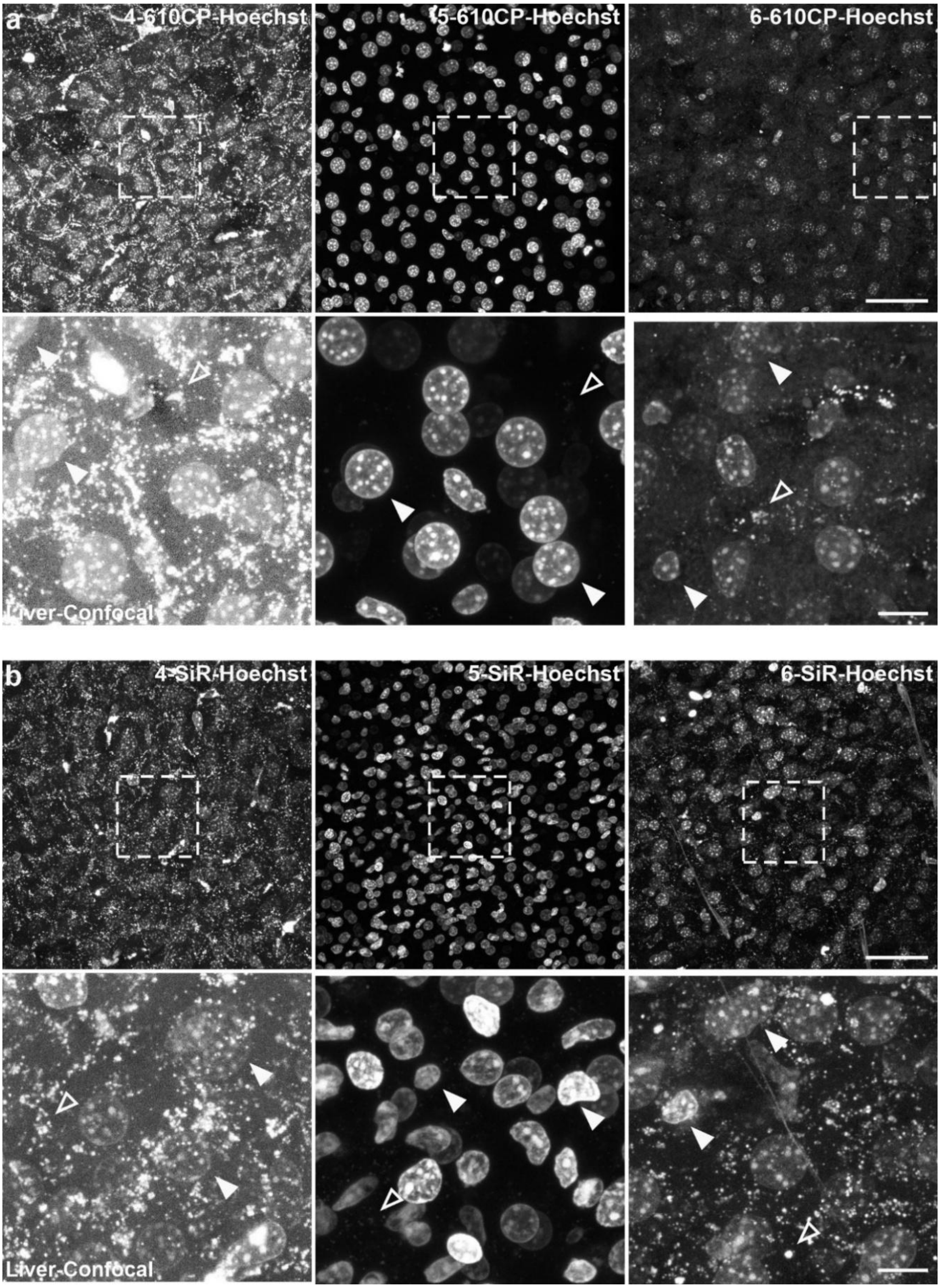

**Figure S13. Staining of nuclei by rhodamine probes in living mouse liver.**

Confocal images of liver living tissue isolated from mice injected prior 24 h with 4-, 5-, 6-610CP-Hoechst (**a**) and 4-, 5-, 6-SiR-Hoechst (**b**). Images were acquired on Leica TCS SP8 confocal system. Images in upper rows are overviews. Images in lower rows show magnified insets of areas delineated in overviews. Full arrowheads point to stained nuclei, empty arrowheads indicate nonspecific staining. Scale bars: 50 µm in overview (upper row), 10 µm – zoomed images (lower row).

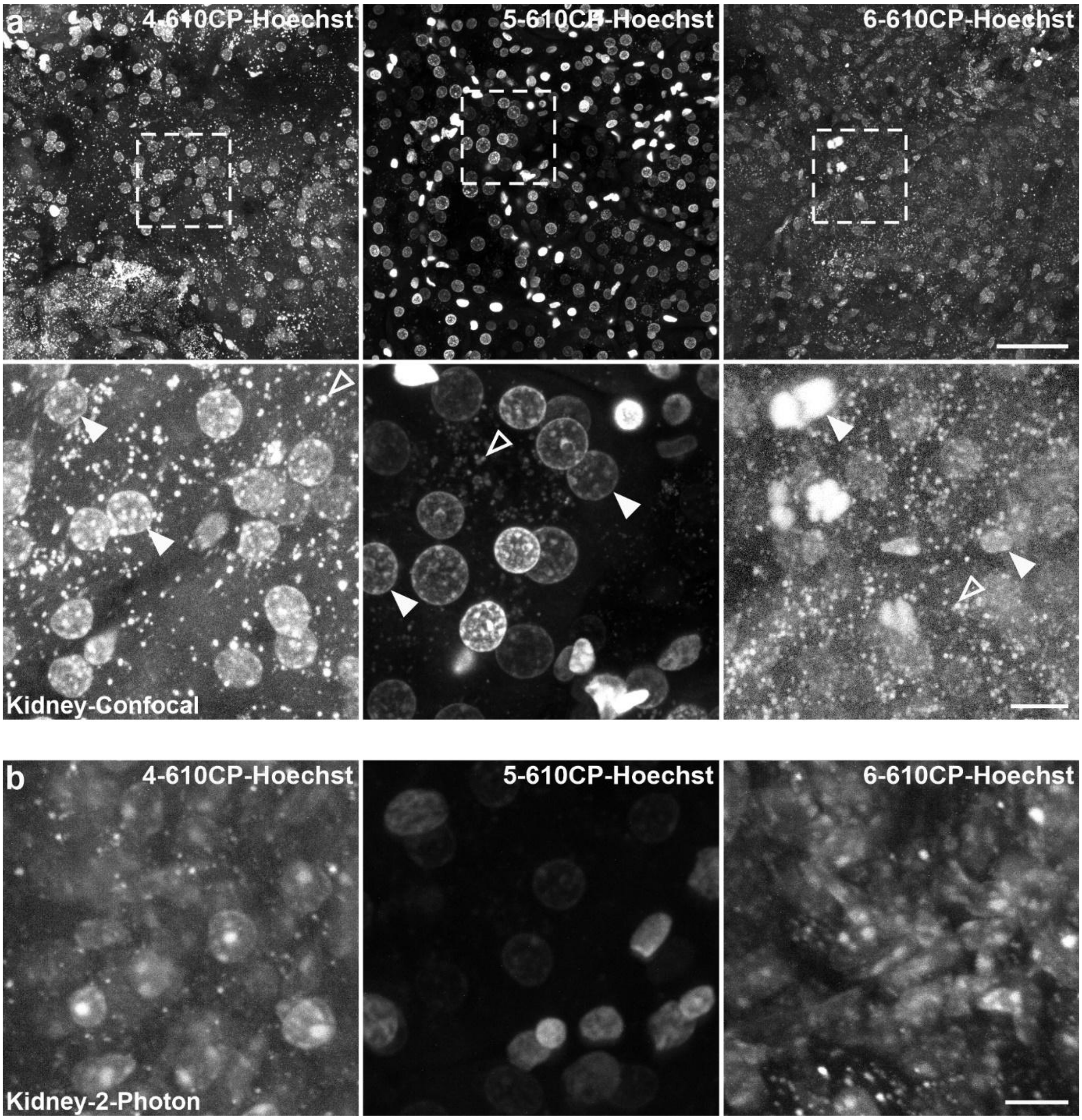

**Figure S14. Staining of nuclei by 610CP-Hoechst probes in living mouse kidney.**

(**a**) Confocal images of kidney living tissue isolated from mice injected prior 24 h with 4-, 5-, 6-610CP-Hoechst. Full arrowheads point to stained nuclei, empty arrowheads indicate nonspecific staining. Scale bars: 50 µm in overview (upper row), 10 µm – zoomed images (lower row). Images were acquired on Leica TCS SP8 confocal system. (**b**) Two-photon microscopy images of kidney living tissues isolated from mice injected prior 24 h with 4-, 5-, 6-610CP-Hoechst. Scale bar: 10 µm. LaVision TriM Scope II multiphoton microscope.

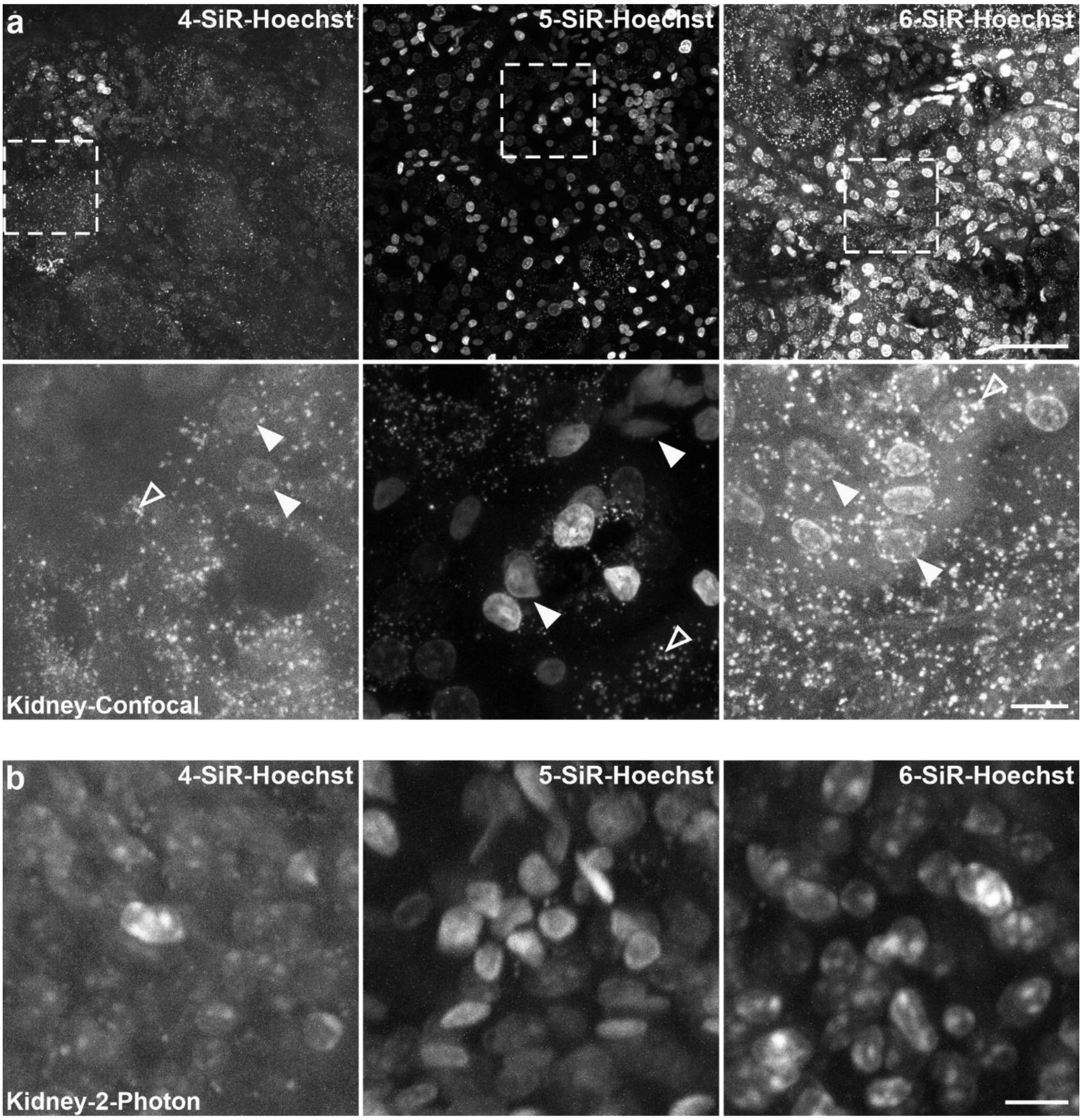

**Figure S15. Staining of nuclei by SiR-Hoechst probes in living mouse kidney.**

(**a**) Confocal images of kidney living tissue isolated from mice injected prior 24 h with 4-, 5-, 6-SiR-Hoechst. Full arrowheads point to stained nuclei, empty arrowheads indicate nonspecific staining. Scale bars: 50 µm in overview (upper row), 10 µm – zoomed images (lower row). Images were acquired on Leica TCS SP8 confocal system. (**b**) Two-photon microscopy images of kidney living tissues isolated from mice injected prior 24 h with 4-, 5-, 6-SiR-Hoechst. Scale bar: 10 µm. LaVision TriM Scope II multiphoton microscope.

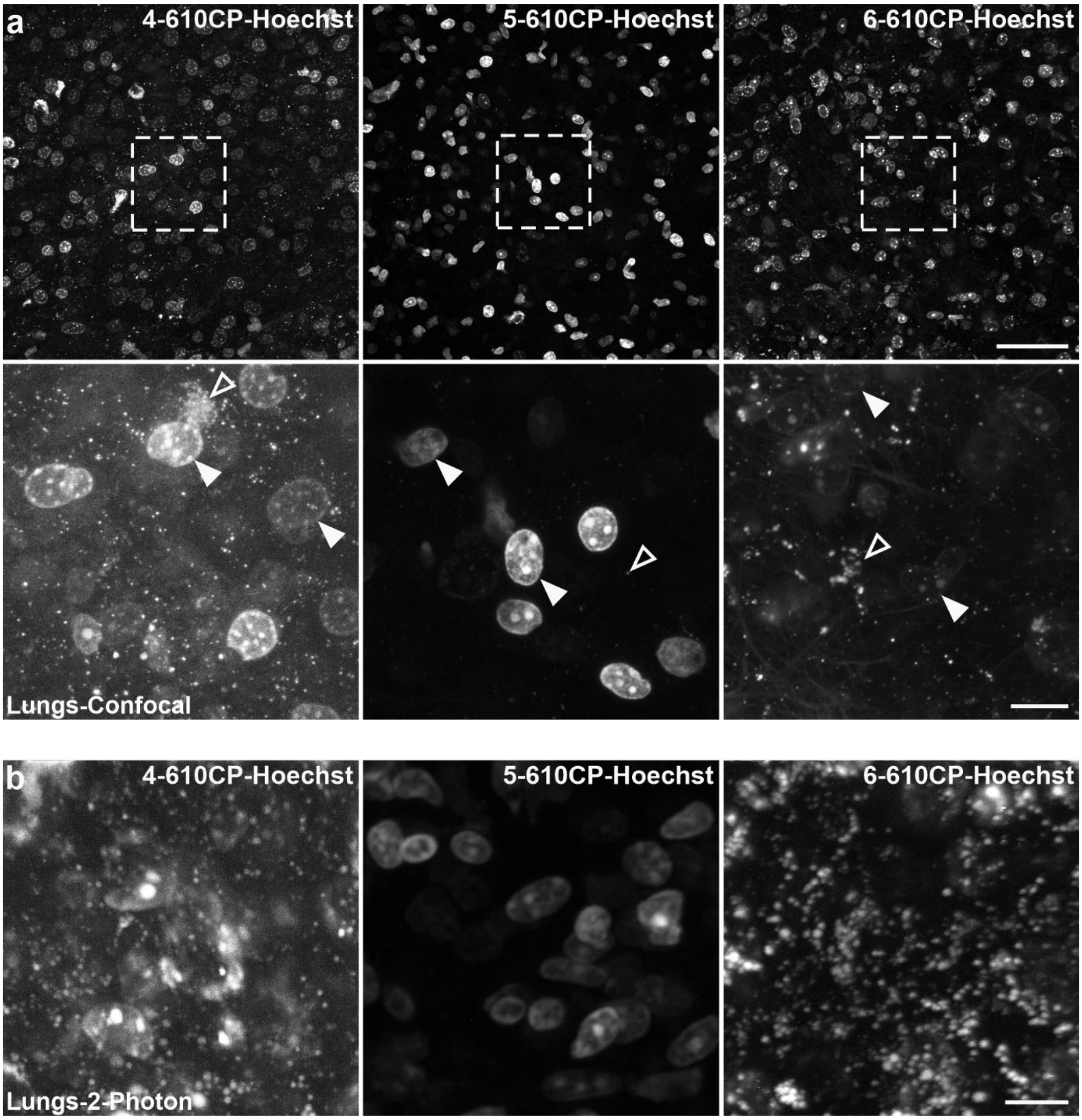

**Figure S16. Staining of nuclei by 610CP-Hoechst probes in living mouse lungs.**

(**a**) Confocal images of lungs living tissue isolated from mice injected prior 24 h with 4-, 5-, 6-610CP-Hoechst. Full arrowheads point to stained nuclei, empty arrowheads indicate nonspecific staining. Scale bars: 50 µm in overview (upper row), 10 µm – zoomed images (lower row). Images were acquired on Leica TCS SP8 confocal system. (**b**) Two-photon microscopy images of kidney living tissues isolated from mice injected prior 24 h with 4-, 5-, 6-610CP-Hoechst. Scale bar: 10 µm. LaVision TriM Scope II multiphoton microscope.

.

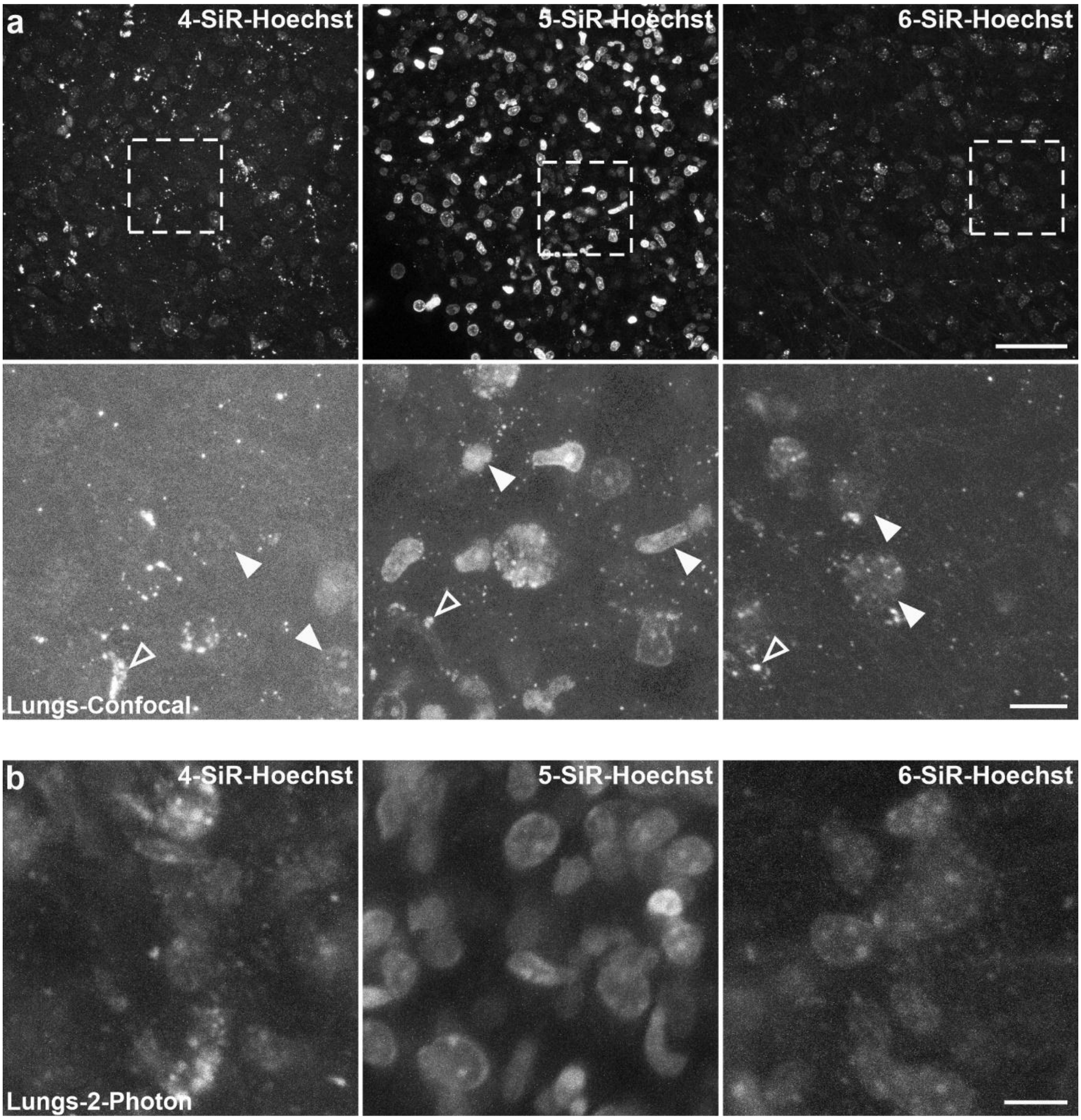

**Figure S17. Staining of nuclei by SiR-Hoechst probes in living mouse lungs.**

(**a**) Confocal images of lungs living tissue isolated from mice injected prior 24 h with 4-, 5-, 6-SiR-Hoechst. Full arrowheads point to stained nuclei, empty arrowheads indicate nonspecific staining. Scale bars: 50 µm in overview (upper row), 10 µm – zoomed images (lower row). Images were acquired on Leica TCS SP8 confocal system. (**b**) Two-photon microscopy images of kidney living tissues isolated from mice injected prior 24 h with 4-, 5-, 6-SiR-Hoechst. Scale bar: 10 µm. LaVision TriM Scope II multiphoton microscope.

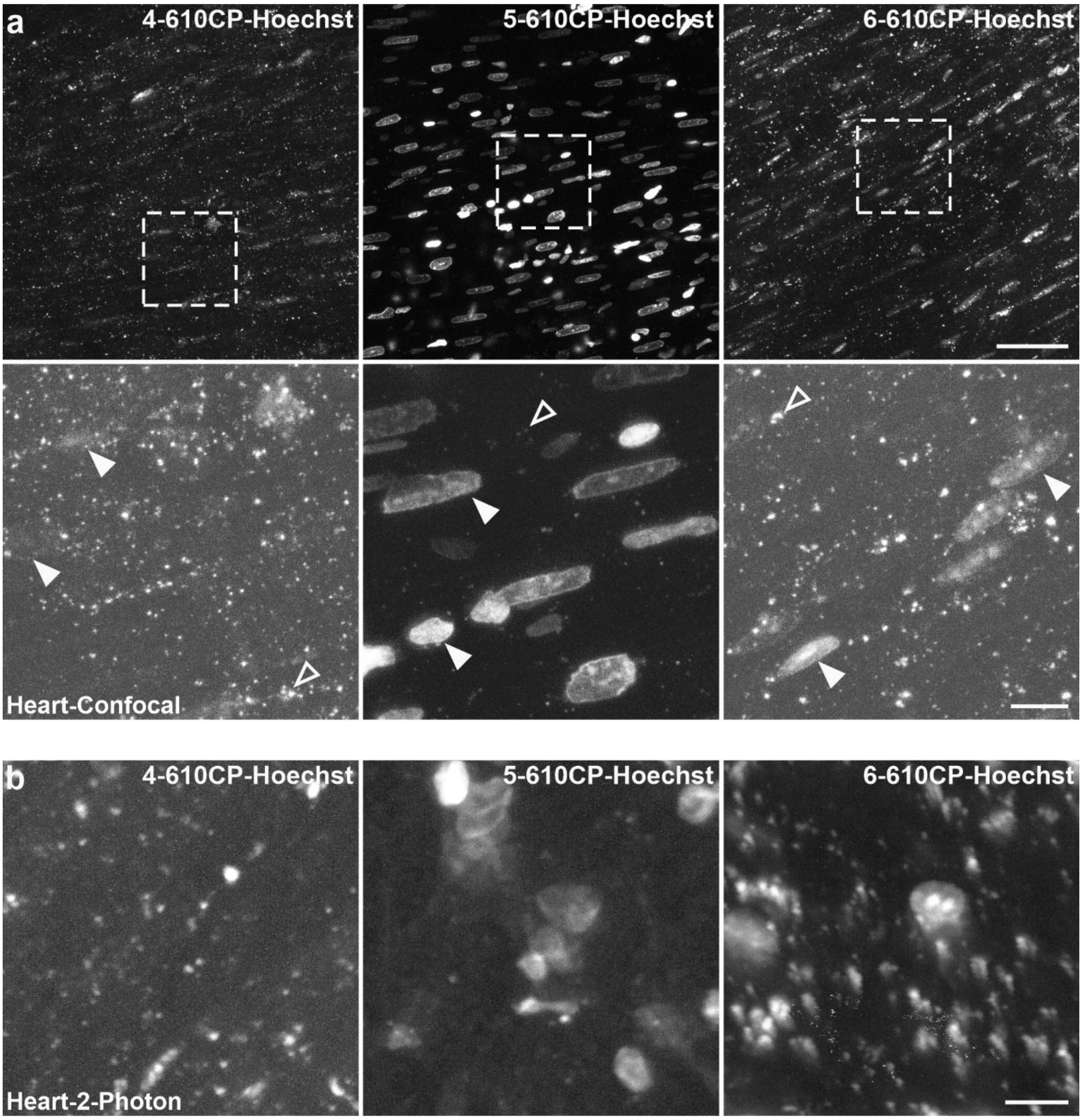

**Figure S18. Staining of nuclei by 610CP-Hoechst probes in living mouse heart.**

(**a**) Confocal images of heart living tissue isolated from mice injected prior 24 h with 4-, 5-, 6-610CP-Hoechst. Full arrowheads point to stained nuclei, empty arrowheads indicate nonspecific staining. Scale bars: 50 µm in overview (upper row), 10 µm – zoomed images (lower row). Images were acquired on Leica TCS SP8 confocal system. (**b**) Two-photon microscopy images of kidney living tissues isolated from mice injected prior 24 h with 4-, 5-, 6-610CP-Hoechst. Scale bar: 10 µm. LaVision TriM Scope II multiphoton microscope.

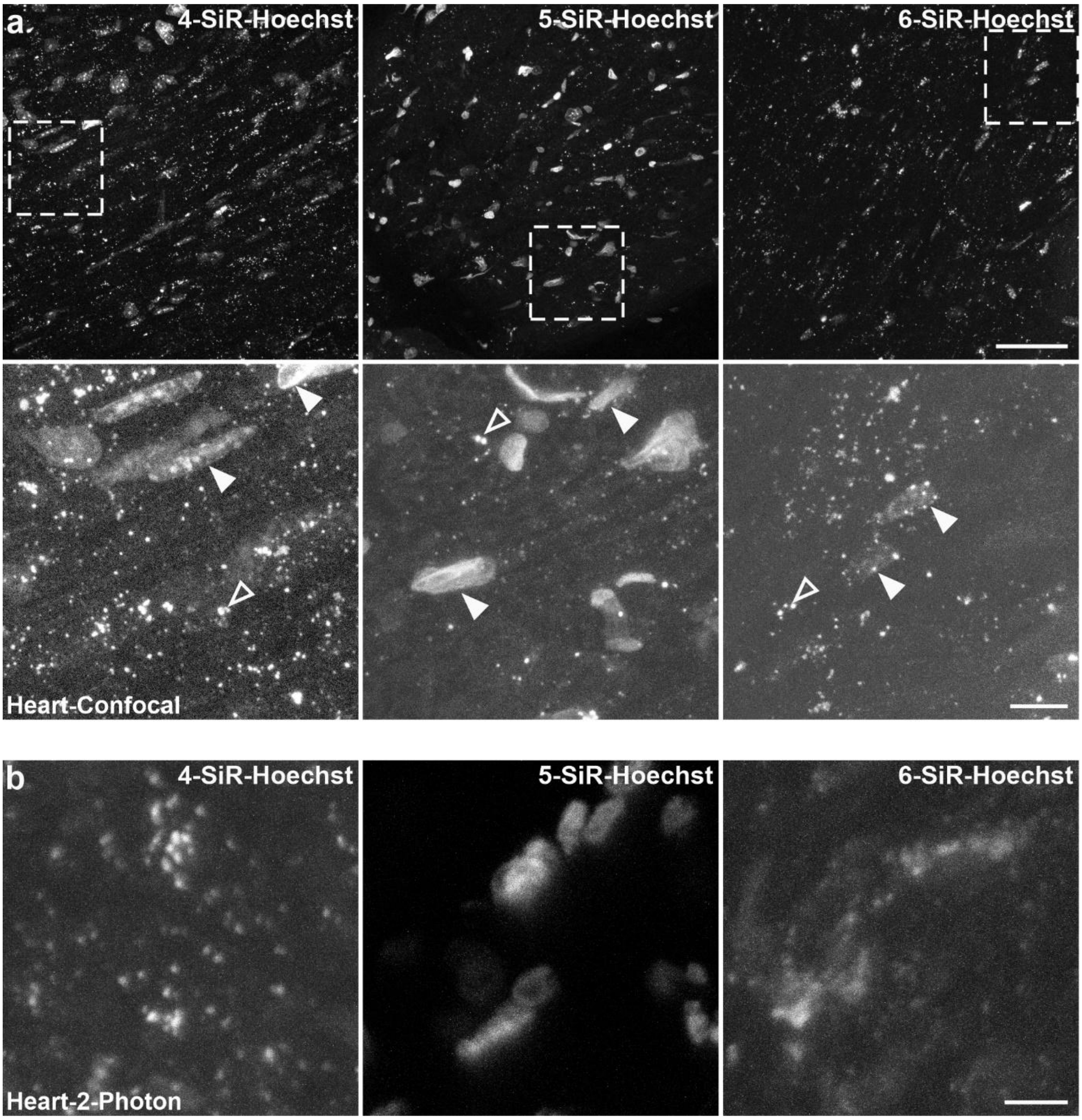

**Figure S19. Staining of nuclei by SiR-Hoechst probes in living mouse heart.**

(**a**) Confocal images of heart living tissue isolated from mice injected prior 24 h with 4-, 5-, 6-SiR-Hoechst. Full arrowheads point to stained nuclei, empty arrowheads indicate nonspecific staining. Scale bars: 50 µm in overview (upper row), 10 µm – zoomed images (lower row). Images were acquired on Leica TCS SP8 confocal system. (**b**) Two-photon microscopy images of kidney living tissues isolated from mice injected prior 24 h with 4-, 5-, 6-SiR-Hoechst. Scale bar: 10 µm. LaVision TriM Scope II multiphoton microscope.

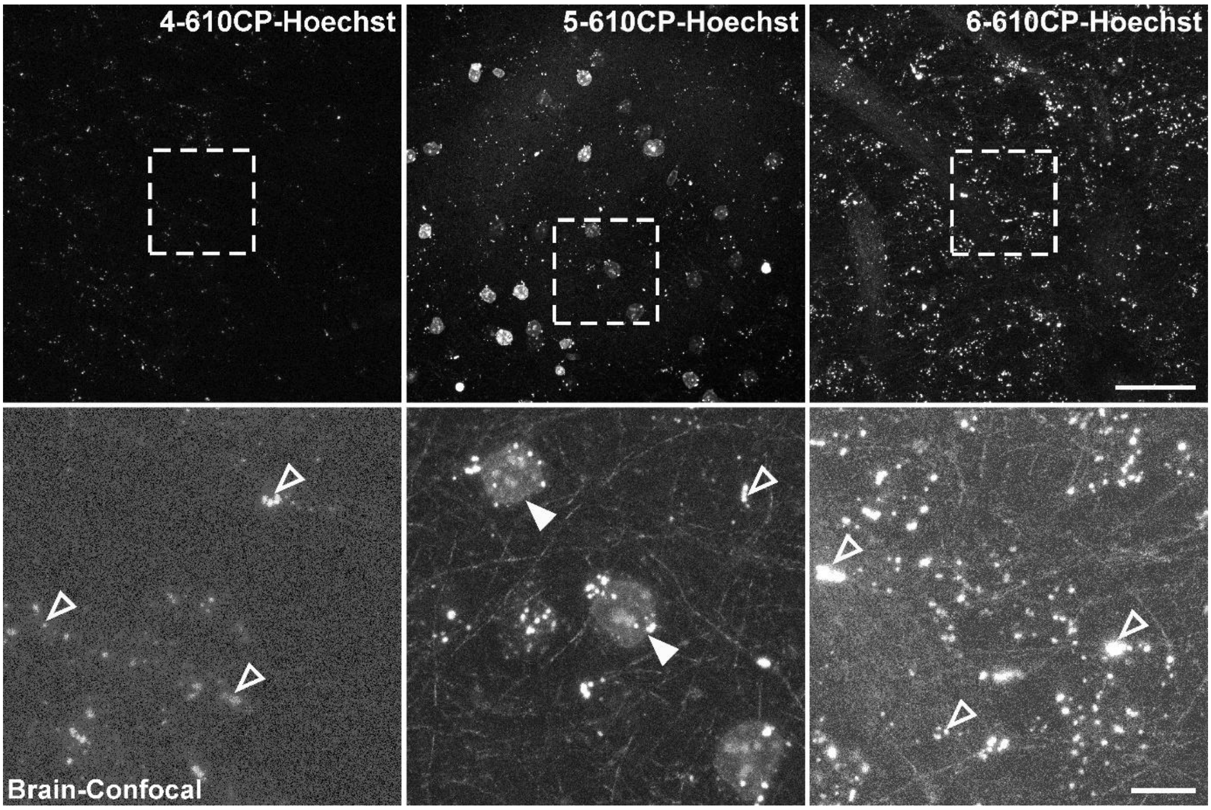

**Figure S20. Staining of nuclei by 610CP-Hoechst probes in living mouse brain.**

Confocal images of brain living tissue isolated from mice injected prior 24 h with 4-, 5-, 6-610CP-Hoechst. Full arrowheads point to stained nuclei, empty arrowheads indicate nonspecific staining. Scale bars: 50 µm in overview (upper row), 10 µm – zoomed images (lower row). Images were acquired on Leica TCS SP8 confocal system.

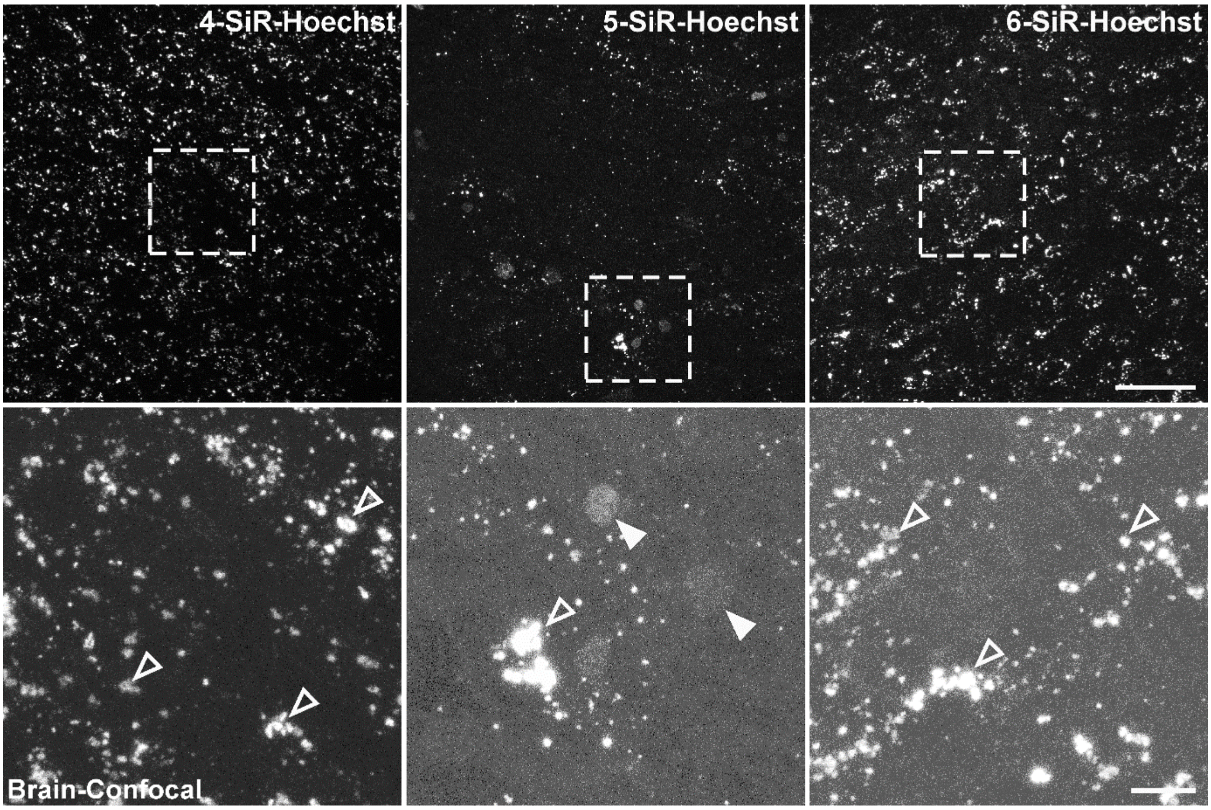

**Figure S21. Staining of nuclei by SiR-Hoechst probes in living mouse brain.**

Confocal images of brain living tissue isolated from mice injected prior 24 h with 4-, 5-, 6-SiR-Hoechst. Full arrowheads point to stained nuclei, empty arrowheads indicate nonspecific staining. Scale bars: 50 µm in overview (upper row), 10 µm – zoomed images (lower row). Images were acquired on Leica TCS SP8 confocal system.

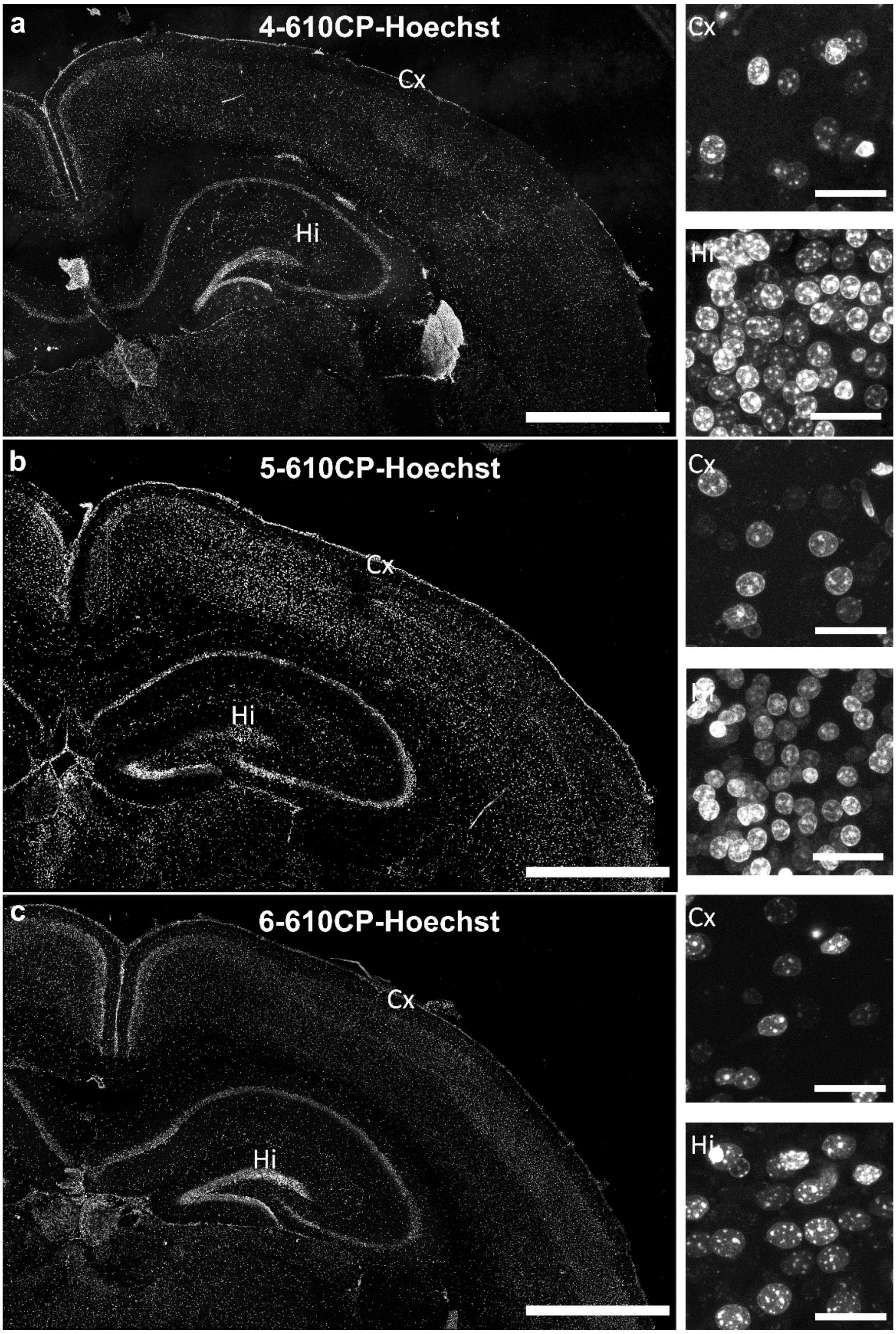

**Figure S22. *Ex-vivo* staining of nuclei by 4-, 5-, 6-610CP-Hoechst probes in mouse coronal brain sections.**

Confocal images of brain tissues isolated from mice and incubated with 250 nM (a) 4-, (b) 5-, (c) 6-610CP-Hoechst probes. Images are acquired on Leica TCS SP8 confocal system. The images on the left are overviews of mouse brain slices stained with the indicated DNA probes. Smaller images on the right show magnified nuclei in cortex (Cx) and hippocampus (Hi) regions. Scale bar: 1 mm for the big overviews on the left and 20 µm for the insets.

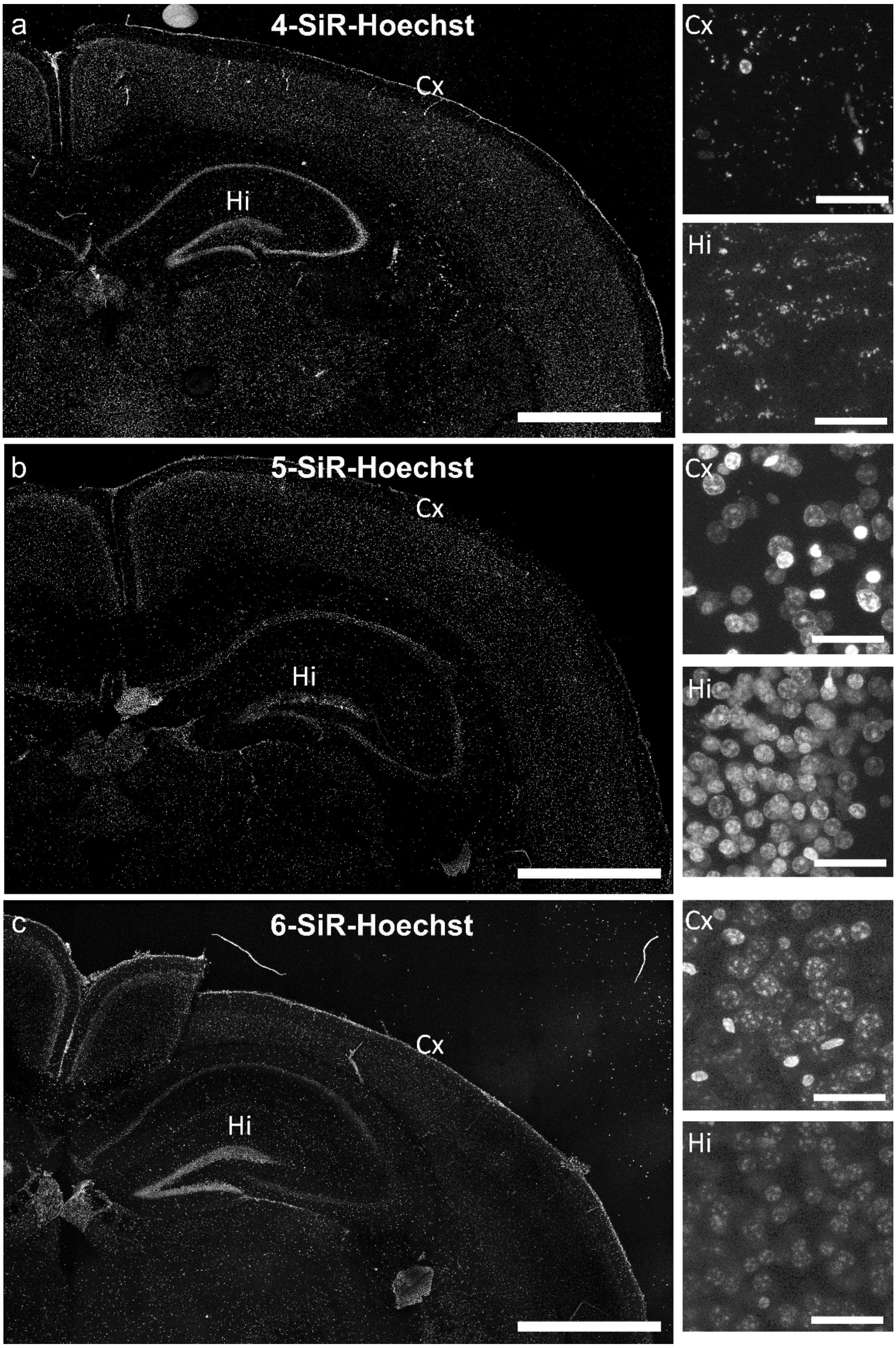

**Figure S23.** ***Ex-vivo*** **staining of nuclei by 4-, 5-, 6-SiR-Hoechst probes in mouse coronal brain sections.**

Confocal images of brain tissues isolated from mice and incubated with 250 nM (a) 4-, (b) 5-, (c) 6-SiR-Hoechst probes. Images are acquired on Leica TCS SP8 confocal system. The images on the left are overviews of mouse brain slices stained with the indicated DNA probes. Smaller images on the right show magnified nuclei in cortex (Cx) and hippocampus (Hi) regions. Scale bar: 1 mm for the big overviews on the left and 20 µm for the insets.

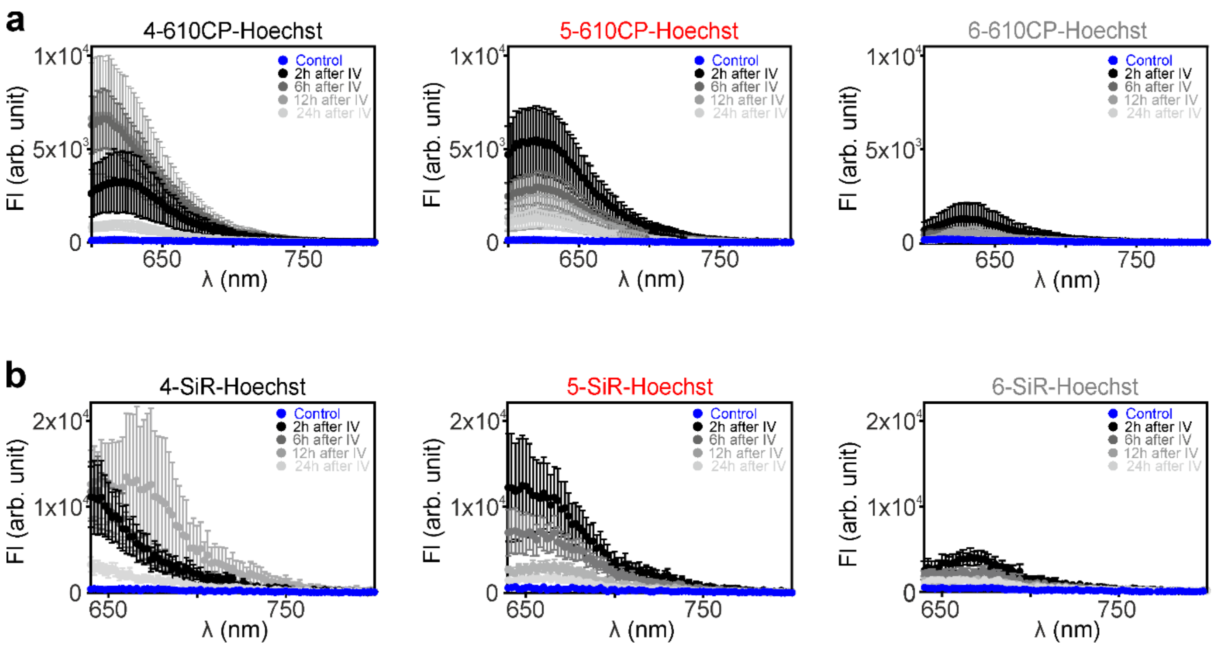

**Figure S24. Excretion of rhodamine-Hoechst probes in urine at different time points.**

The fluorescence intensity of urine samples measured at 2, 6,12 and 24 h after injection of 610CP-Hoechst (**a**) or SiR-Hoechst (**b**) probes. Experimental data are presented as mean ± SEM of 3≤n≤5 independent experiments.
